# Exercise Related microRNAs in *C. elegans* Regulate Calcium Homeostasis and Mitochondrial Dynamics: Conserved pathways, Divergent mircoRNAs

**DOI:** 10.1101/2025.05.13.653755

**Authors:** Qin Xia, Jiwang Tang, José C. Casas-Martinez, Penglin Li, Leo R Quinlan, Katarzyna Goljanek-Whysall, Brian McDonagh

## Abstract

Exercise can help mitigate age-related muscle atrophy, promoting mitochondrial function, Ca^2+^ homeostasis and regulating gene expression. MicroRNAs (miRs) are crucial post-transcriptional regulators of gene expression, fine tuning protein levels to maintain cellular homeostasis. In *C. elegans*, a 5-day swimming regimen increased mitochondrial content, lifespan and fitness. Small RNA sequencing identified exercise specific miRs, including increased levels of cel-miR-57-5p and cel-miR-249-3p, and decreased cel-miR-72-3p and cel-miR-77-5p. *mir-57* and *mir-249* mutant strains had enhanced fertility, survival and lifespan, whereas *mir-72* and *mir-77* mutant strains had diminished fertility, survival and lifespan. Interestingly both *miR-77* and *mir-249* mutant strains had disrupted mitochondrial morphology and reduced fitness following exercise. The exercise related miRs identified in *C. elegans* did not have conserved mammalian orthologs. Exercise regulated mammalian miRs were identified from the literature including mmu-miR-181a-5p, mmu-miR-199a-5p and mmu-miR-378a-3p. Treatment of murine myoblasts with mmu-miR-181a-5p and mmu-miR-378a-3p enhanced mitochondrial content, autophagy markers and myogenesis, while mmu-miR-199a-5p impaired these processes. Exercise related miRs identified in *C. elegans* target genes regulating Ca^2+^ homeostasis such as *ipp-5* (inositol Polyphosphate-5-phosphatase), *sca-1* (Ca^2+^ transporting ATPase) and *ncx-2* (mitochondrial Na^+^/Ca^2+^ antiporter). It was also confirmed that mmu-miR181a-5p (miR181) targets Inpp5a and antogmiR-181 (AM181) treatment decreased Ca^2+^ handling in myotubes. Similarly, mmu-miR-378a-3p and cel-miR-249-3p target Kinase Suppressor of Ras (Ksr1)/*ksr-2*, involved in the MAPK pathway. Despite direct conservation of exercise related miRs from nematodes to mammals there are common regulatory pathways, contributing to exercise-induced adaptations.

**Graphical abstract:** The exercise related microRNAs (cel-miR-77-5p and cel-miR-249-3p) identified in *C. elegans* following a 5-day swimming protocol are not conserved in mammalians. However, the microRNAs identified share common gene targets regulating Ca^2+^ homeostasis and mitochondrial function, with previously identified exercise related microRNAs in murine models. The results highlight conserved regulatory adaptive mechanisms in response to exercise.

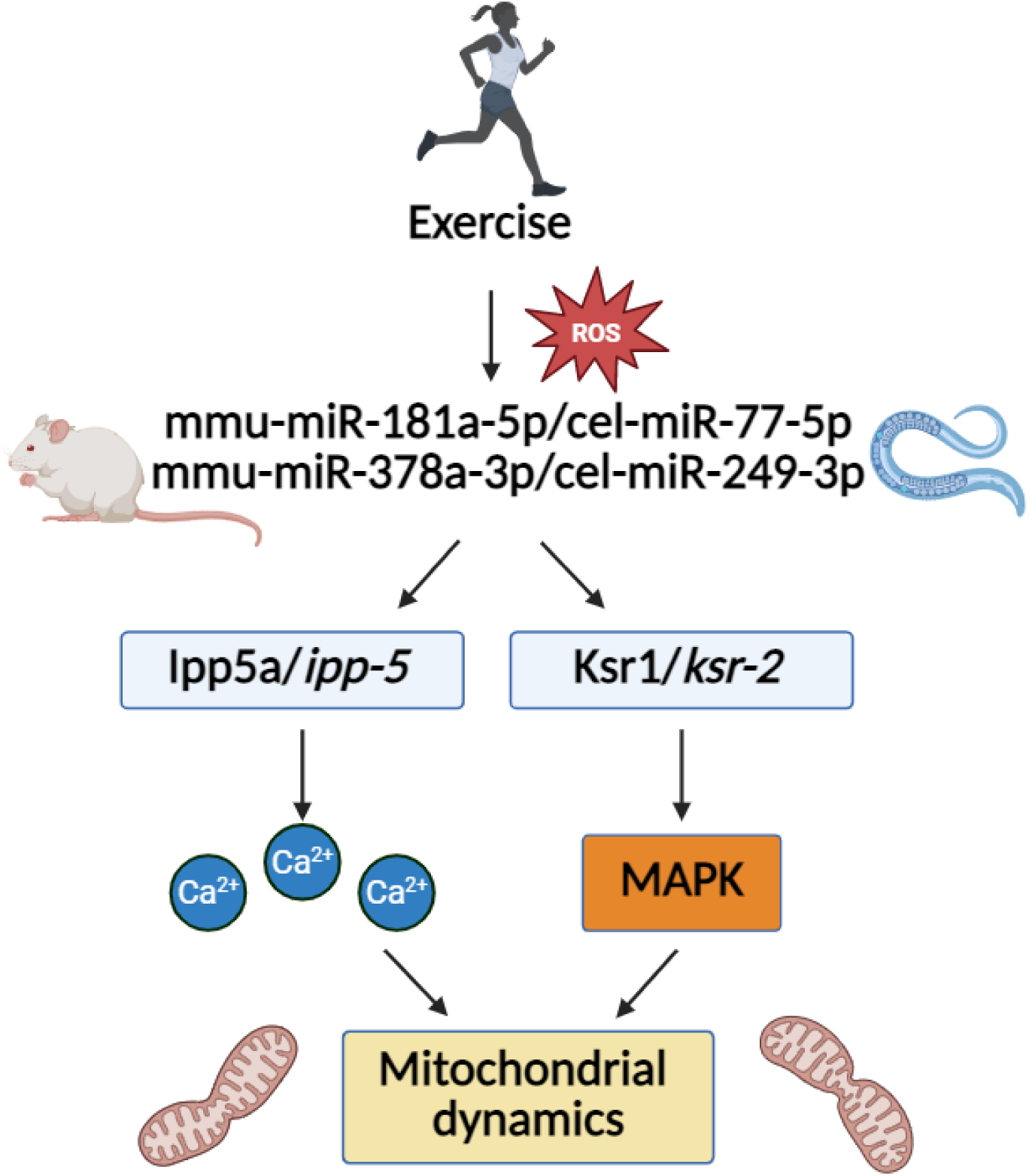

## 1. Introduction

Skeletal muscle is a dynamic and adaptable tissue that can respond to exercise by promoting muscle hypertrophy and metabolic efficiency [1], whereas disuse, ageing or disease can result in muscle atrophy [2]. Regular exercise can help maintain muscle mass but also exerts multifaceted beneficial effects on age-related diseases, including coronary atherosclerosis, osteoarthritis and sarcopenia [3–5]. Age-related muscle atrophy presents significant public health challenges due to its association with elevated rates of hospitalisation, loss of independence and mortality [6]. Exercise is an effective approach for maintaining muscle mass and function [7]. Nevertheless, the mechanisms underlying the adaptive response to exercise are yet to be fully understood. Mitochondrial dysfunction is considered a hallmark of ageing and increased mitochondrial dysfunction has been described in skeletal muscle with age [8, 9]. Exercise has been demonstrated to promote mitochondrial biogenesis and dynamics in skeletal muscle with enhanced metabolic efficiency [10, 11]. During contractile activity there is endogenous generation of reactive oxygen species (ROS) [12], resulting in an acute stress response and activation of key signalling pathways required for the beneficial adaptive response to exercise [13]. Redox sensitive transcription factors such as NRF2, NF-κB and FOXO are activated in response to an altered intracellular redox environment and activated in response to exercise [14, 15], affecting gene transcription and downstream physiological effects. Redox dependent signalling and post translational modifications can directly affect proteins involved in excitation contraction coupling and Ca^2+^ handling [16] but also regulate key signalling pathways involved in mitochondrial dynamics, proteostasis and muscle repair [17, 18].

In response to exercise the acute activation of redox sensitive transcription factors can directly affect the transcriptional response. However, the adaptive response to exercise can be fine-tuned at the post transcriptional level by changes in levels of distinct microRNAs (miRs) [19, 20]. miRs are small non-coding RNAs approximately 19–25 nucleotides in length with the first 2-8 nucleotides containing the seed sequence that typically binds to the 3’UTR of target genes [21]. miRs play a pivotal role in the modulation of post-transcriptional gene expression and are reported to regulate 2/3 of the human genome [22]. miRs can inhibit translation of mRNAs and promote mRNA degradation by loading into RNA induced silencing complexes [22, 23]. Furthermore, one miR can potentially target a range of different genes often within the same signalling pathway, and individual genes can be regulated by distinct miRs, further adding complexity to the system. A number of exercise specific miRs have been identified in a range of models from humans to rodents [20]. Many of these miRs contain binding sites for exercise related transcription factors such as NRF2, NF-κB, FOXO etc but also target these redox dependent transcription factors providing a negative feedback loop to regulate antioxidant response, mitochondrial dynamics and Ca^2+^ handling. Potentially this provides a mechanism to fine tune the adaptive response to exercise.

*Caenorhabditis elegans* is an established and powerful whole animal model to investigate cellular signalling pathways. The body wall muscle of *C. elegans* closely resembles the sarcomere of striated muscle in vertebrates and is a valuable model for investigation of muscle maintenance and function [24]. Many of the key intracellular signalling pathways are conserved in *C. elegans* and it has been utilised as a model to determine the adaptive response to exercise [25]. It has been demonstrated that an exercise intervention can increase lifespan, help maintain body wall muscle, promote physical activity and delay the onset of neurodegeneration [26, 27]. Following exercise in *C. elegans*, an acute increase in ROS generation has been demonstrated, including activation of SKN-1 (NRF2 ortholog) and DAF-16 (FOXO ortholog), resulting in the promotion of mitochondrial turnover and content [28, 29]. Moreover, it was demonstrated that exercise promotes Mitochondrial ER Contact Sites (MERCS) assembly and mitochondrial remodelling [28]. However, the effects of exercise on miR levels in *C. elegans* has not previously been investigated.

In this study, a 5-day exercise programme that has previously been demonstrated to increase mitochondrial content and improved healthspan was performed in *C. elegans* [27]. Small RNA sequencing of miRs, revealed changes in specific miRs following exercise. Specifically, there were increased levels of cel-miR-57-5p and cel-miR-249-3p, whereas cel-miR-72-3p and cel-miR-77-5p decreased in response to exercise. Furthermore *mir-57* and *mir-249* mutant strains had enhanced fertility, survival and lifespan, whereas *mir-72* and *mir-77* mutant strains had diminished fertility, survival and lifespan. Interestingly both *miR-77* and *mir-249* mutant strains exhibited impaired mitochondrial morphology and reduced fitness following exercise. cel-miR-77-5p targets a number of genes regulating Ca^2+^ homeostasis such as *ipp-5* (inositol Polyphosphate-5-phosphatase), *sca-1* (Ca^2+^ transporting ATPase) and *ncx-2* (mitochondrial Na^+^/Ca^2+^ antiporter). Although the exercise related miRs identified in *C. elegans* were not conserved in mammals. Exercise related mammalian miRs identified from the literature: mmu-miR-181a-5p, mmu-miR-199a-5p and mmu-miR-378a-3p, shared targets with the exercise related miRs identified in *C. elegans*. The levels of mmu-miR-181a-5p, mmu-miR-199a-5p and mmu-miR-378a-3p increased in response to an acute physiological bolus of H_2_O_2_ in an *in vitro* myoblast model. Treatment of myoblasts with mmu-miR-181a-5p and mmu-miR-378a-3p mimics led to enhanced mitochondrial content, autophagy markers and improved myogenesis. Among the shared target genes and pathways were the regulation of Ca^2+^ homeostasis and mitochondrial associated genes. Specifically, the results demonstrate that mmu-miR-181a-5p and cel-miR-77-5p commonly target Inositol Polyphosphate-5-Phosphatase A (Inpp5a)/*ipp-5* that reduces inositol 3 phosphate levels (IP_3_), contributing to the regulation of Ca^2+^ metabolism in muscle. Ca^2+^ imaging was performed in myotubes following treatment with synthetic miR-181a (miR181) and AntagomiR-181a (AM181), revealing decreased Ca^2+^ flux with AM181 and elevated Inpp5a. Furthermore, mmu-miR-378a-3p and cel-miR-249-3p were found to share a common target Kinase Suppressor of Ras (Ksr1)/*ksr-2*, activating the Mitogen activated protein kinase (MAPK) pathway involved in mitochondrial biogenesis. The inter-species conservation in exercise related miRs targeting provides insights into potential conserved regulatory mechanisms in cellular processes related to Ca^2+^ homeostasis and mitochondrial function.

## 2. Materials and methods

### 2.1 *C. elegans* strains and swimming exercise

*C. elegans* were cultured on nematode growth medium (NGM) plates seeded with *E. Coli* (OP50) at 20 °C. *C. elegans* stains N2 wild type, MT16311 (*mir-77(n4286) II*), MT13015 (*mir-72(n4130) II*), MT16848 (*mir-249(n4983) X*), VC347 (*mir-57(gk175) II*) and SJ4103 (*zcIs14 [myo-3::gfp(mit)]*) strains were obtained from the Caenorhabditis Genetics Center (CGC). The outcrossing of *mir-77* and *mir-249* strains with mitochondrial muscle GFP reporter SJ4103 *zcIs14 [myo-3::GFP(mit)]* were performed in this project. The 5-day swimming protocol was performed as previously described [27].

### 2.2 C2C12 cell culture

C2C12 myoblasts were cultured in 10% growth medium supplemented with DMEM, 10% fetal bovine serum and 1% penicillin/streptomycin. Cells were maintained under standard conditions at 37 ℃ with 5% CO_2_. To investigate the impact of cholesterol-linked miR-181, 199, 378 mimic and antagomiR-181, 199, 378, C2C12 myoblasts at 50-60% confluency was subjected to treatment with a final concentration of 100 nM for 24 hours prior to an acute 10-minute exposure to 25 µM H_2_O_2_, that has previously been demonstrated to improve myogenesis and increased mitochondrial content [27]. Following treatment, media was exchanged and cells were allowed to recover for 24 hours before protein and RNA were extracted. In order to assess the impact of these miRs on myotube formation, C2C12 cells at 70-80% confluence were treated with the miRs and antogmiRs (100 nM) for 24 hours prior to H_2_O_2_ treatment, growth media were replaced with differentiation media and the cells were cultured in differentiation media for a period of 5-7 days. MF20 staining was performed to determine effects on myogenesis.

### 2.3 Western blotting

Proteins were quantified using the Bradford assay, 20 µg of protein were separated on 12% SDS-PAGE gels and transferred using a semi-dry blotter. After blocking with 5% milk, the membranes were incubated overnight with primary antibodies (rabbit anti-TOM20, VDAC1, SIRT1, ATG5, ULK1, SQSTM1/p62 and LC3b) at a 1:1000 dilution. Following washing with TBS-T, the membranes were incubated with secondary antibodies (IRDye 800CW Goat anti-Mouse IgG) at a 1:10,000 dilution in the dark for 1 hour. The acquired images of the blot were captured using the Odyssey Fc imaging system. Ponceau S staining was used to confirm total protein loading for normalisation. The quantification and normalisation of the blots were analysed utilising Image Studio Lite.

### 2.4 PCR for genotyping

DNA extraction from *C. elegans* was performed using worm lysis mix (WLM) containing proteinase K. 15 worms were added to a PCR tube along with 15ul of WLM and placed at -80 °C for 1 hour. Subsequently samples were transferred to a thermocycler and subjected to incubation at 65°C for 1 hour, followed by a denaturation step at 95°C for 5 minutes and then cooled on ice. The PCR reaction mix was prepared by combining 12.5 μl of MyTaq™ Reaction Buffer Red, 0.5 μl each of forward and reverse primers, 9.0 μl of nuclear-free water and 2.5 μl of DNA. The amplification process commenced with an initial denaturation step at 95 °C for 10 minutes, followed by 40 cycles consisting of denaturation at 95 °C for 15 seconds, annealing at 56 °C for 15 seconds and extension at 72 °C for 90 seconds. Finally, the reaction was held at 4 °C. Subsequently, the PCR products were subjected to agarose electrophoresis and gel images were captured using the LI-COR Odyssey.

### 2.5 qPCR

RNA extraction from both cellular and *C. elegans* samples was performed using TRIzol [30]. For mRNA synthesis, 500 ng of RNA was mixed with 1 μl of random hexamers and incubated at 65 °C for 10 minutes. Then, the mixture was combined with a master mix including 4 μl of RT buffer, 2 μl of DTT, 1 μl of dNTP, 1 μl of Superscript II and 1 μl of Ribolock, followed by incubation at 42 °C for 60 minutes. For miR synthesis, a mixture was prepared by combining 2 μl of 5x miRCury RT reaction buffer, 4.5 μl of RNase-free water, 1 μl of miRCury RT Enzyme mix, 0.5 μl of synthetic RNA spike-in and 2 μl of 5 ng RNA. The mixture was then incubated at 42 °C for 60 minutes, followed by a final incubation at 95 °C for 5 minutes. Subsequent qPCR analysis was conducted using the SYBR Green Master Mix in a 10 µl reaction volume. S29 was used for normalisation of mRNA in cells and miRs were normalised to unisp6 or U6; RNA from worms were normalised to CDC42, both were calculated using the delta–delta Ct method.

### 2.6 microRNA sequencing in *C. elegans*

RNA isolation from *C. elegans* was performed using TRIzol reagent [30]. Briefly, worms in N2 and *prdx-2* mutant strains with or without exercise were dislodged from plates and after the removal of bacteria, TRIzol was employed to disrupt the cuticle of the worms. Subsequently, chloroform was added, followed by centrifugation to isolate the aqueous phase. The final steps involved the addition of isopropanol and ethanol, followed by thorough washing to obtain high-purity RNA. Subsequently, the RNA samples were analysed for small RNA sequencing by TamiRNA (TAmiRNA GmbH Leberstrasse 20 1110 Vienna, Austria) and the assessment of next-generation sequencing (NGS) data quality was conducted through automated and manual evaluation using fastQC v0.11.9 [31] and multiQC v1.10 [32]. Reads from samples underwent adapter trimming and quality filtering with cutadapt v3.3 [33], ensuring a minimum length of 17 nucleotides. Mapping procedures were executed using bowtie v1.3.0 [34] and miRDeep2 v2.0.1.2 [35]. Initially, reads were mapped against the genomic reference WBcel.235 from Ensembl [36], allowing for two mismatches. Subsequently, mapping against miRBase v22.1 was performed [37], filtering for cel-specific miRs and allowing for one mismatch. Statistical analysis of pre-processed NGS data utilised R v4.0 and packages including heatmap vNA, pca Methods v1.82 and genefilter v1.72. Differential expression analysis was carried out using edgeR v3.32 [38], employing quasi-likelihood negative binomial generalised log-linear model functions. To optimise the false discovery rate (FDR) correction, the independent filtering method of DESeq2 was adapted within edgeR [39], specifically for the removal of low-abundance miRs.

### 2.7 Lifespan assay

The lifespan assay was conducted on adult D6 worms following the completion of a chronic exercise regimen, which was performed twice daily for a total of 5 days [27]. A total of 110-120 worms were carefully transferred to new plate every 2 days until all worms died. Survival was assessed daily until the last surviving worm exhibited no response.

### 2.8 Brood size

For the brood size assay, 12 worms at L4 stage worms were transferred to new plates daily. The counting of eggs laid was conducted every 24 hours until the cessation of egg-laying. To determine the egg-laying rate, worms were allowed to lay eggs over a 5-hour period [40].

### 2.9 Oxidative stress assays

In the paraquat and arsenite stress assay, 60 worms, post-chronic exercise, were picked in individual wells of a 96-well plate, each well containing either 100mM Paraquat or 2.5mM Arsenite [41]. Worm death was ascertained by the lack of response to a gentle tap with a picker and survival was assessed every 1-2 hour.

### 2.10 *C. elegans* microscopy

For the MitoTracker staining in *C. elegans*, 40-45 worms were incubated with 2.5 µM MitoTracker Red for 10 minutes in the dark one day after the chronic exercise. Following incubation, worms were then transferred to new NGM plates to remove excess staining in the gut. Subsequently, the worms were immobilised using 20 mM levamisole and imaged at 10× magnification. The fluorescence of the entire body and body length of each worm was measured by ImageJ [27].

To assess mitochondria in the body wall muscle, the SJ4103 strain (*zcIs14 [myo-3::gfp(mit)]*) was utilised and exposed to chronic swimming. After the exercise, worms were immobilised on an agar slide covered with a coverslip. Subsequently, imaging of mitochondria in the body wall muscles, specifically the area between the pharynx and vulva, was performed at 60x magnification. Quantification was performed on 120-150 images and the analysis involved categorising them into five classes based on the criteria outlined in [26].

### 2.11 C2C12 Immunostaining

For MitoTracker Green and MitoSOX staining in cells, medium was removed and washed with Hank’s Balanced Salt Solution (HBSS), followed by incubation with a 200 nM MitoTracker Green solution for 30 minutes. The cells were incubated with a 5 µM MitoSOX Red solution for 10 minutes. After each incubation step, the cells were washed with HBSS. Following staining, images were captured at 60x magnification. The identification of mitochondrial areas within each cell from various fields of view was based on the presence of MitoTracker Green. Subsequently, the fluorescence intensity within Regions of Interest (ROIs) containing either MitoTracker Green or MitoSOX staining was quantified using ImageJ software as previously described [27].

For MF20 staining of myotubes, media was removed and fixed in ice cod methanol for 5 minutes, then blocked in 10% horse serum for 1 hour at room temperature on rocker, followed by incubation in MF20 primary antibody for overnight. On the subsequent day, the primary antibody was removed and incubated with second antibody for 1 hour, followed by 5 minutes of DAPI staining and the images were captured.

### 2.12 CeleST

To evaluate the physiological activity of *C. elegans*, CeleST analysis was performed the following methodology outlined in prior research [42]. Briefly, 4-5 worms (with a total of at least 30 worms) were recorded for 30 seconds using a Nikon LV-TV microscope set at 1× magnification and the physiological activity quantified using CeleST [42].

### 2.13 miR target prediction and validation

The predicted targets of the selected miRs were compared to see if common pathways were targeted by exercise related miRs in *C. elegans* and mammals. The predicted targets of exercise-related miRs, cel-miR-77-5p and cel-miR-249-3p in *C. elegans*, mmu-miR-181a-5p, mmu-miR-199a-5p and mmu-miR-378a-3p in mammalian cells, were determined using TargetScan Release 7.1 (https://www.targetscan.org/mmu_71/). In *C. elegans*, there were 145 predicted targets of cel-miR-77-5p, 103 predicted targets of cel-miR-249-3p. In C2C12, there were 1371 predicted targets of mmu-miR-181a-5p, 634 predicted targets of mmu-miR-199a-5p, 223 predicted targets of mmu-miR-378a-3p. Bioinformatic analysis identified conserved targets of cel-miR-77-5p and cel-miR-249-3p in *C. elegans*, as well as mmu-miR-181a-5p, mmu-miR-199a-5p and mmu-miR-378a-3p in mice. qPCR was employed to validate a subset of these putative shared targets across the two species. TransmiR v2.0 database (https://www.cuilab.cn/transmir) was used to determine miRs regulated by exercise-related TFs (NRF2, NF-κB, FOXO3, STAT3).

### 2.14 Ca^2+^ imaging in myotubes

Myotubes growing on glass coverslips were loaded with Fluo-4 AM (2.5 μM) in complete medium for 15 minutes at 37°C, followed by a 15-minute wash period in dye-free complete medium. Coverslips were transferred to a recording chamber filled with physiological buffer (135 mM NaCl, 5 mM KCl, 1 mM MgCl₂, 2 mM CaCl₂, 10 mM HEPES, 10 mM glucose, pH 7.45) and imaged using a ZEISS Axiovert 200M inverted microscope equipped with a xenon lamp (LAMBDA LS, excitation range: 330–650 nm) and Smart Shutter Lambda 10/3 controller. Fluorescence images were captured at 1 Hz for 5 minutes. Excitation and emission wavelengths were set to 488 nm and 510 nm (long-pass filter), respectively. Baseline fluorescence was recorded for 1–2 minutes. At 2 minutes, ATP (100 μM final concentration) was added, followed by KCl (50 mM final concentration) at 4 minutes. For inhibition experiments, cells were pre-incubated with the purinergic receptor antagonist suramin (200 μM) for 2 minutes prior to recording. Image stacks were analysed using the SARFIA plugin [43] in Igor Pro 8. Regions of interest (ROIs) were defined based on predefined fluorescence thresholds. Data were normalised using the formula ΔF/F0, where F0 represents the average baseline fluorescence over the initial 10 seconds. Peak responses, mean, area under curve, and full-width half-maximum (FWHM) fluorescence values were extracted using the NeuroMatic plugin in Igor Pro 8 [44].

### 2.15 Statistics

Images were analysed semi-automatically using ImageJ and Image Studio Lite, with subsequent manual corrections. Western blot band intensities were normalised to protein levels determined by Ponceau S staining. For C2C12 microscopy, 3–6 images per biological replicate were randomly captured at 10× or 40× magnification from different fields of view. Myogenic differentiation was assessed via MF20 immunostaining, measuring myotube diameter as the average width of all myotubes per field of view, myotube area as the average area fraction, and fusion index as the percentage of nuclear within myotubes relative to the total nuclear. For the MitoTracker and MitoSOX staining were quantified based on relative fluorescence intensity per cell. For SJ4103 strain staining, 130–150 images were analysed in the region between the pharynx and vulva, classified into five predefined categories. Statistical analyses are detailed in the respective figure legends. Differences between two groups were assessed using a student t-test, while one-way or two-way ANOVA was applied for comparisons involving multiple groups. The log-rank test was used for oxidative stress tolerance assays, and chi-square tests were performed for categorical distributions. A *p* value < 0.05 was considered statistically significant. GraphPad Prism 7 was used for all statistical analyses and graphical representations. The source data supporting figure generation are available in the corresponding source data file.

## 3 Results

### 3.1 cel-miR-57-5p and cel-miR-249-3p levels increased but cel-miR-72-3p and cel-miR-77-5p levels decreased following chronic swimming exercise

To identify changes in miRs expression in *C. elegans* in response to exercise, small RNA sequencing was performed following a 5-day swimming exercise regimen. The exercise protocol has previously been shown to enhance mitochondrial capacity, improve physical fitness and extend both survival and longevity in wild-type worms [27]. The exercise regimen resulted in the upregulation of 5 miRs and the downregulation of 17 miRs in the N2 wild type strain (Fig. 1a). The *prdx-2* mutant strain has a disrupted response to exercise [27, 28], 14 miRs were upregulated and 15 miRs were downregulated in this strain (Fig. 1b). A comparative analysis between the control groups of N2 and *prdx-2* mutant worms revealed 3 upregulated and 5 downregulated miRs (Fig. S1). Detailed fold changes and corresponding *p*-values for these miRs are provided in Table S1. Interestingly a previous proteomic analysis in the *prdx-2* mutant strain identified disrupted microRNA processing [27], following exercise miR-37, miR-39 and miR-72 exhibited opposing changes in the levels of -3p and -5p mature miRs, suggesting arm switching and miR biogenesis defects (Table S1). In the N2 wild type and *prdx-2* mutant stain cel-miR-57-5p and cel-miR-249-3p were upregulated, whereas cel-miR-72-3p and cel-miR-77-5p were downregulated following exercise suggesting their physiological relevance to exercise in *C. elegans*.

**Figure 1.**
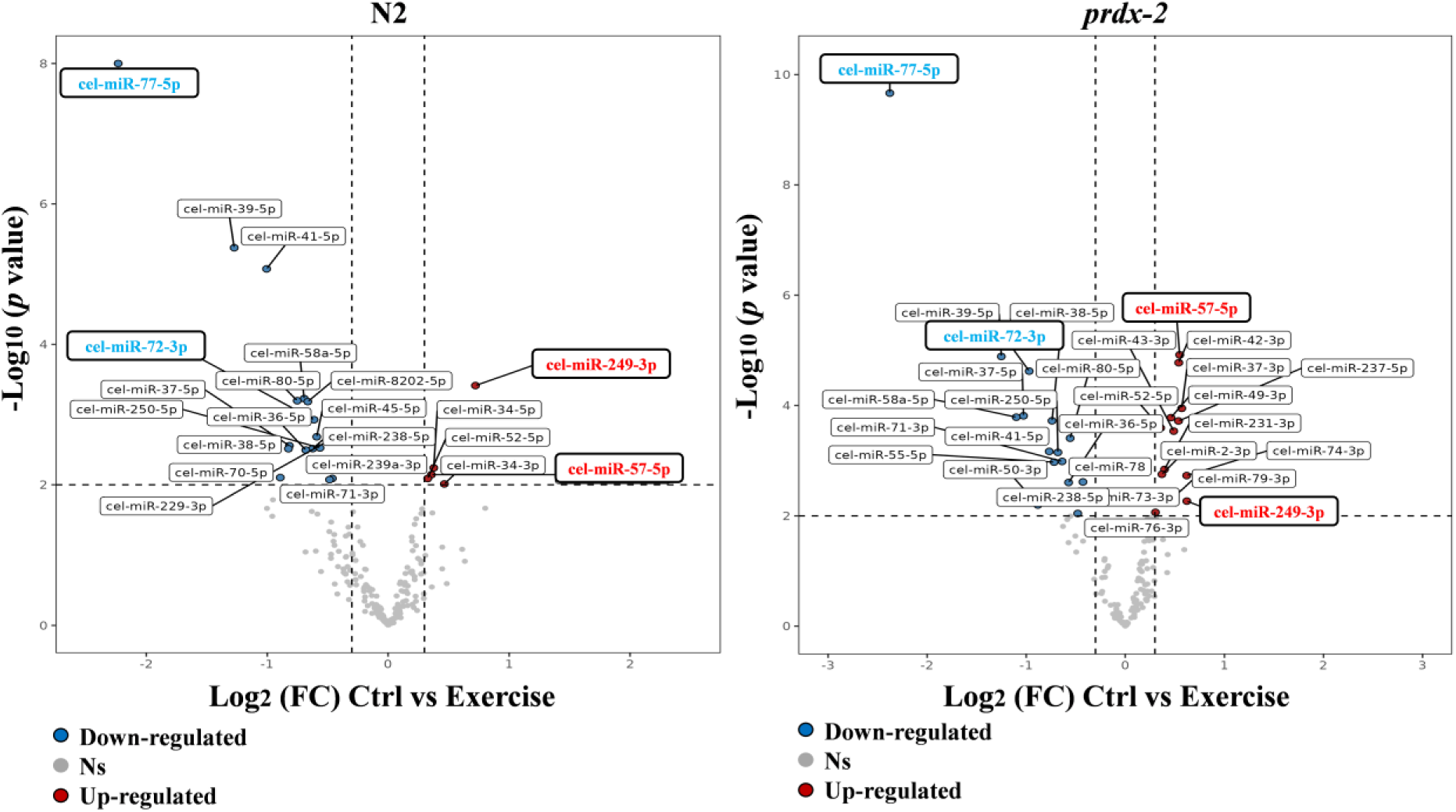
Differential miR expression in *C. elegans* following 5-day swimming protocol. Comparison of miR expression between control and exercised groups in N2 wild type FDR 9.0% (a); Comparison of miR expression between control and exercised groups in the *prdx-2* mutant FDR 4.5% (b). Data represent 4 independent replicates per condition. A total of 378 miRs were identified and quantified. miRs with an adjusted *p*-value < 0.01 and a minimum fold change > 1.5 were considered significantly altered by exercise.

### 3.2 cel-miR-57-5p, cel-miR-72-3p, cel-miR-77-5p and cel-miR-249-3p regulate fertility, survival and lifespan in *C. elegans*

To investigate the functional roles of cel-miR-57-5p, cel-miR-72-3p, cel-miR-77-5p and cel-miR-249-3p, we used N2 wild type and mutant strains of common miRs upregulated (cel-mir-57-5p and cel-mir-249-3p) and downregulated (cel-mir-72-3p and cel-mir-77-5p) following exercise. Brood size and egg laying rates of the strains were assessed. The *mir-57* and *mir-249* mutant strains had decreased brood size and egg laying rate compared to N2 wild type, while the *mir-72* and *mir-77* mutant strains had increased brood size (Fig. 2a-c). Subsequently, we performed lifespan and oxidative stress assays including paraquat and arsenite exposure following the swimming exercise. In the N2 control non-exercised group, the *mir-57* and *mir-249* mutants exhibited an increased lifespan and enhanced resistance to paraquat compared to N2 wild type, while the *mir-72* and *mir-77* mutants demonstrated a reduced lifespan and decreased resistance to both paraquat and arsenite (Fig. 2d-f). In the exercised groups, lifespan and survival increased in the N2 wild type strain, but decreased in the *mir-57*, *mir-72* and *mir-249* mutants (Fig. S2). The exact mean lifespan and corresponding *p* values for these mutants can be found in Table S2. These findings underscore the critical roles of cel-miR-57-5p, cel-miR-72-3p, cel-miR-77-5p and cel-miR-249-3p in modulating fertility, survival and lifespan, highlighting their importance in the physiological response to exercise.

**Figure 2.**
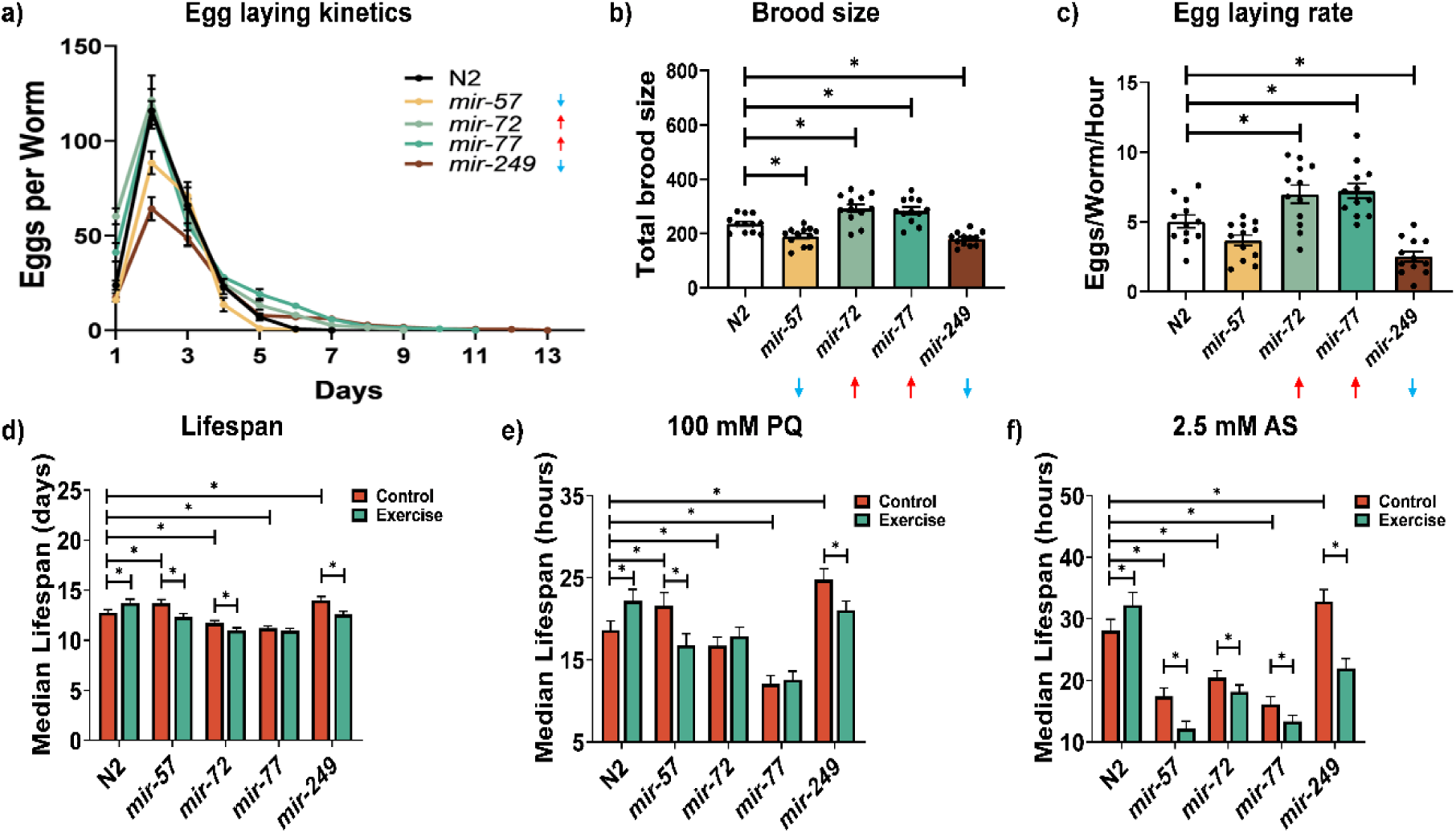
cel-miR-57-5p, cel-miR-72-3p, cel-miR-77-5p and cel-miR-249-3p regulate fertility, survival and lifespan in *C. elegans*. cel-miR-57-5p, cel-miR-72-3p, cel-miR-77-5p and cel-miR-249-3p regulate fecundity (a-c), lifespan (d) and resistance to oxidative stress following exercise at day 6 (e-f). Graphs are the normalised relative means ± SEM and all experiments were performed with n = 12 (egg laying assay), n=110-120 (lifespan assay), n=60 (oxidative stress assay) and *p*-value of < 0.05 was considered as statistically significant *(p < 0.05). *p* values (b: N2 vs *mir-57* = 0.0187, N2 vs *mir-72* = 0.0026, N2 vs *mir-77* = 0.0133, N2 vs *mir-249* = 0.0034. c: N2 vs *mir-72* = 0.0233, N2 vs *mir-77* = 0.01, N2 vs *mir-249* = 0.0023. d: N2 vs *mir-57* = 0.0467, N2 vs *mir-72* = 0.0103, N2 vs *mir-77* = 0.0002, N2 vs *mir-249* = 0.0105; N2: Control vs Exercise = 0.0432; *mir-57*: Control vs Exercise = 0.0039; *mir-72*: Control vs Exercise = 0.0364; *mir-249*: Control vs Exercise = 0.0019. e: N2 vs *mir-57* = 0.0362, N2 vs *mir-77* < 0.0001, N2 vs *mir-249* = 0.0003; N2: Control vs Exercise = 0.0228; *mir-57*: Control vs Exercise = 0.0144; *mir-249*: Control vs Exercise = 0.0097. f: N2 vs *mir-57* < 0.0001, N2 vs *mir-72* = 0.0002, N2 vs *mir-77* < 0.0001, N2 vs *mir-249* = 0.0305; N2: Control vs Exercise = 0.0453; *mir-57*: Control vs Exercise = 0.0114; *mir-72*: Control vs Exercise = 0.0402; *mir-77*: Control vs Exercise = 0.0369; *mir-249*: Control vs Exercise < 0.0001)

### 3.3 cel-miR-57-5p, cel-miR-72-3p, cel-miR-77-5p and cel-miR-249-3p influence mitochondrial potential and morphology in *C. elegans*

Given the established link between lifespan, survival ability and mitochondrial function [45], the impact of these miRs on mitochondrial membrane potential and morphology following the swimming exercise was determined. MitoTracker Red staining was used as an indicator of mitochondrial membrane potential. In the non-exercised control groups, the *mir-249* mutant strain increased staining intensity compared to N2 wild type while the *mir-57*, *mir-72* and *mir-77* mutants had decreased staining (Fig. 3a). In the exercise group, MitoTracker intensity was enhanced in N2 wild type following exercise but decreased in the *mir-57*, *mir-72* and *mir-249* mutant strains. Given the consistent observations in the *mir-77* and *mir-249* mutant strains, the mutants were crossed with the muscle mitochondrial reporter strain SJ4103 *zcIs14 [myo-3p::mitogfp]*. Mitochondrial morphology was quantified based on five distinct categories, each representing a progressive increase in mitochondrial fragmentation and disorganisation [25]. Consistent with the MitoTracker Red staining, the N2 and *mir-249* mutant strains, contained an abundance of interconnected mitochondria in the body wall muscle, whereas the *mir-77* mutant exhibited increased mitochondrial fragmentation in the body wall muscle (Fig. 3b). Additionally, the swimming exercise increased the filamentous mitochondria in N2 wild type strain but resulted in increased mitochondrial fragmentation in both *mir-77* and *mir-249* mutants.

**Figure 3.**
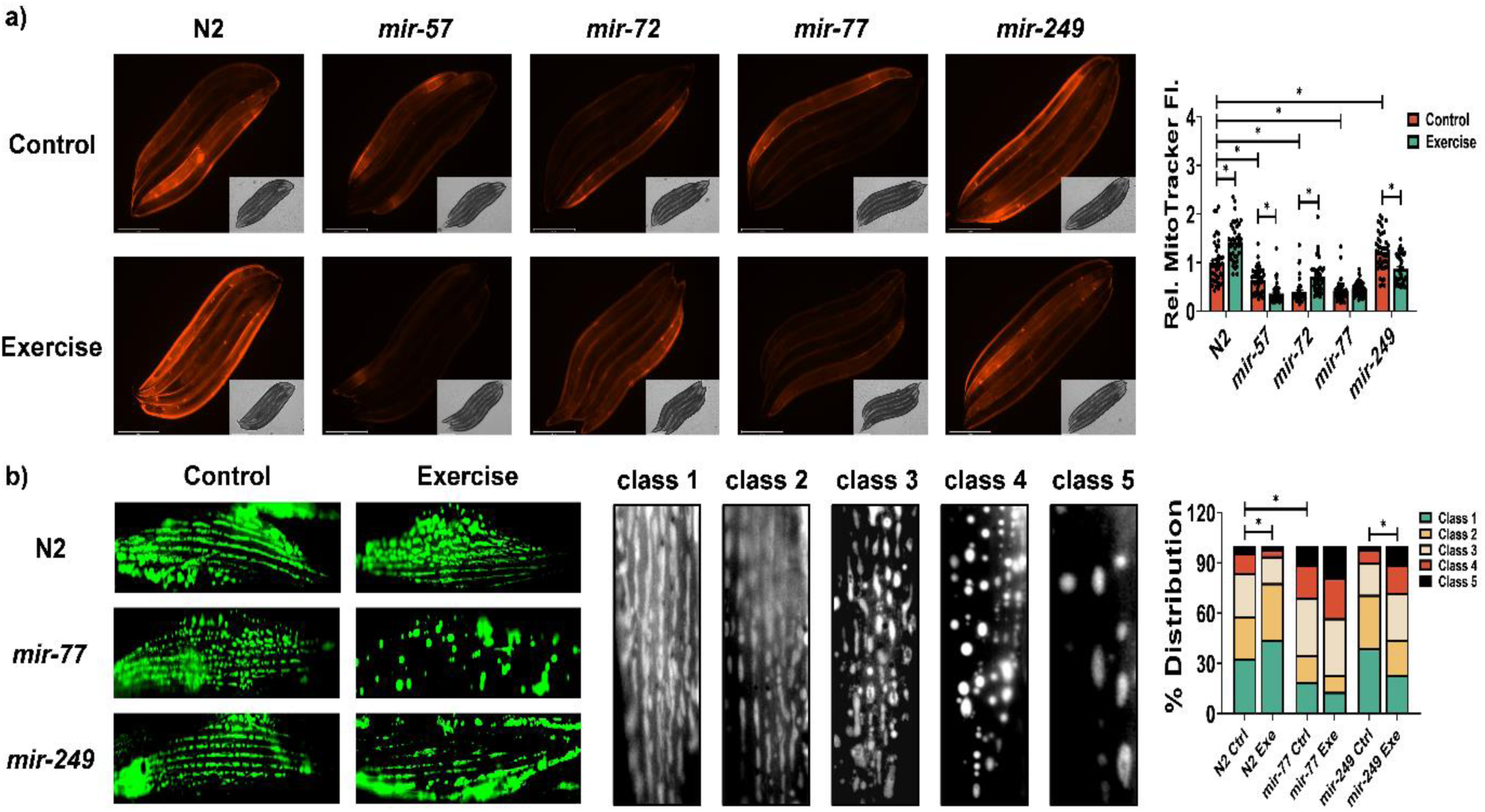
cel-miR-77-5p and cel-miR-249-3p are required for improved mitochondrial capacity following swimming exercise in *C. elegans*. Representative images of MitoTracker red staining for mitochondrial membrane potential (scale bar = 275 μm) and *myo-3::gfp* reporter strain for mitochondrial morphology (scale bar = 50 μm) following swimming exercise at day 6 (a-b). Graphs are the normalised relative means ± SEM and all experiments were performed with n = 40-45 (MitoTracker red staining), n=130-150 (mitochondria reporter) and *p*-value of < 0.05 was considered as statistically significant *(p < 0.05). *p* values (a: MitoTracker: N2 vs *mir-57* < 0.0001, N2 vs *mir-72* < 0.0001, N2 vs *mir-77* < 0.0001, N2 vs *mir-249* = 0.0299; N2: Control vs Exercise < 0.0001; *mir-57*: Control vs Exercise = 0.0011; *mir-72*: Control vs Exercise < 0.0001; *mir-249*: Control vs Exercise < 0.0001. b: N2 vs *mir-77* = 0.0168; N2: Control vs Exercise = 0.0406; *mir-249*: Control vs Exercise = 0.0015)

### 3.4 cel-miR-57-5p, cel-miR-72-3p, cel-miR-77-5p and cel-miR-249-3p influence physical fitness in *C. elegans*

To evaluate the roles of these exercise related miRs on physical activity, CeleST was used to track the swimming activity [42]. Key metrics such as activity index, wave initiation rate, travel speed and brush stroke were quantified as indicators of enhanced physical fitness, while body wave number, asymmetry, stretch and curling were assessed as markers of frailty [42]. In the non-exercised control groups, the *mir-77* mutant strain exhibited a decreased activity index and wave initiation rate compared to N2 wild type (Fig. 4a,b). Similarly, the *mir-72* displayed a reduced activity index along with increased stretch (Fig. 4a,d). Following exercise, fitness parameters such as activity index, wave initiation rate and travel speed increased in the N2 wild-type (Fig. 4a-b, S3a). However, the *mir-249* mutant showed a decreased wave initiation rate and activity index, while no significant changes were observed in the other three mutants.

**Figure 4.**
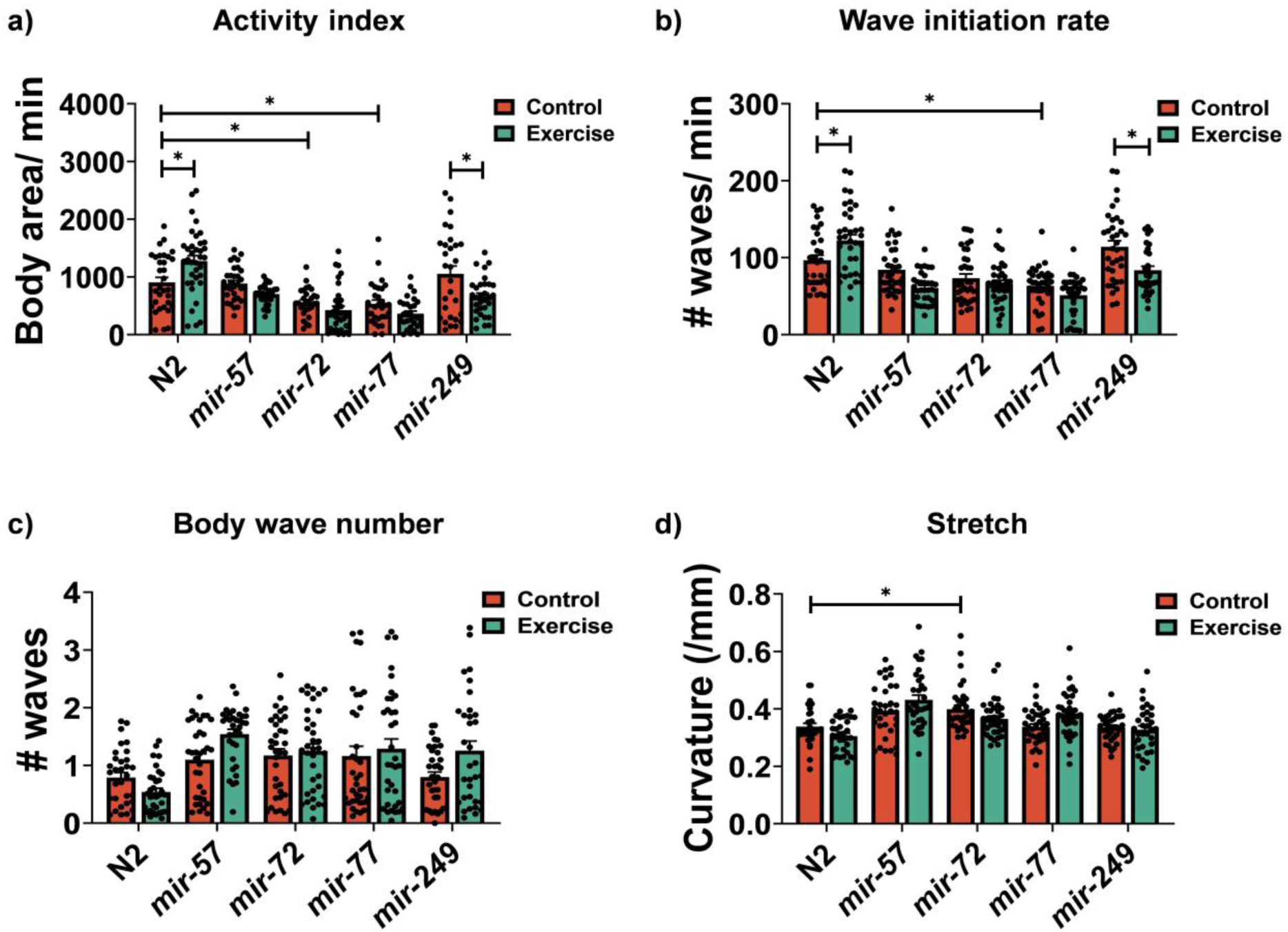
cel-miR-77-5p and cel-miR-249-3p are required for physical fitness following swimming exercise in *C. elegans*. Activity index (a), Wave initiation rate (b), body wave number (c) and stretch (d) following chronic swimming exercise at day 6 (a-h). Graphs are the normalised relative means ± SEM and all experiments were performed with n = 30-40 and *p*-value of < 0.05 was considered as statistically significant *(p < 0.05). *p* values (a: N2 vs *mir-72* = 0.0461, N2 vs *mir-77* = 0.0179; N2: Control vs Exercise = 0.0112; *mir-249*: Control vs Exercise = 0.0109. b: N2 vs *mir-77* = 0.0013; N2: Control vs Exercise = 0.042. d: N2 vs *mir-72* = 0.0222)

### 3.5 mmu-miR-181a-5p, mmu-miR-199a-5p and mmu-miR-378a-3p are up-regulated following the treatment of physiological levels of H_2_O_2_

A number of exercise related miRs have been identified in mammals regulating a range of physiological functions from mitochondrial turnover to Ca^2+^ homeostasis. When we compared the seed sequences of the miRs identified in *C. elegans* that are responsive to exercise with their mammalian miRs, no conserved miRs or seed sequences were identified, highlighting that not all miRs exhibit evolutionary conservation across species [46]. Previous work demonstrated an increase in the nuclear localisation of redox-sensitive transcription factors (NRF2, NF-κB, FOXO3a and STAT3) following brief exposure to physiological levels of H_2_O_2_ [27]. The TransmiR v3.0 database (http://www.cuilab.cn/transmir) was used to identify exercise-related miRs that contain binding sites for these transcription factors including mmu-miR-181a-5p, mmu-miR-199a-5p and mmu-miR-378a-3p (Table 1). NRF2 and STAT3 jointly regulate miR-181 [47, 48], NF-κB regulates miR-199 and miR-378 [49–51]. Moreover, these miRs have been previously shown to play crucial roles in the regulatory processes of skeletal muscle myogenesis [52] and in response to exercise (Table 1). miR-181a-5p levels increase with exercise and decrease with age [53, 54], it has been reported to regulate key components of mitochondrial turnover and enhances myogenesis through the regulation of Hox-a11 [55]. miR-199 inhibits myogenesis by targeting WNT signaling pathway [56]. miR-378 regulates myogenesis by targeting MyoR and enhancing MyoD expression [57].

**Table 1.**
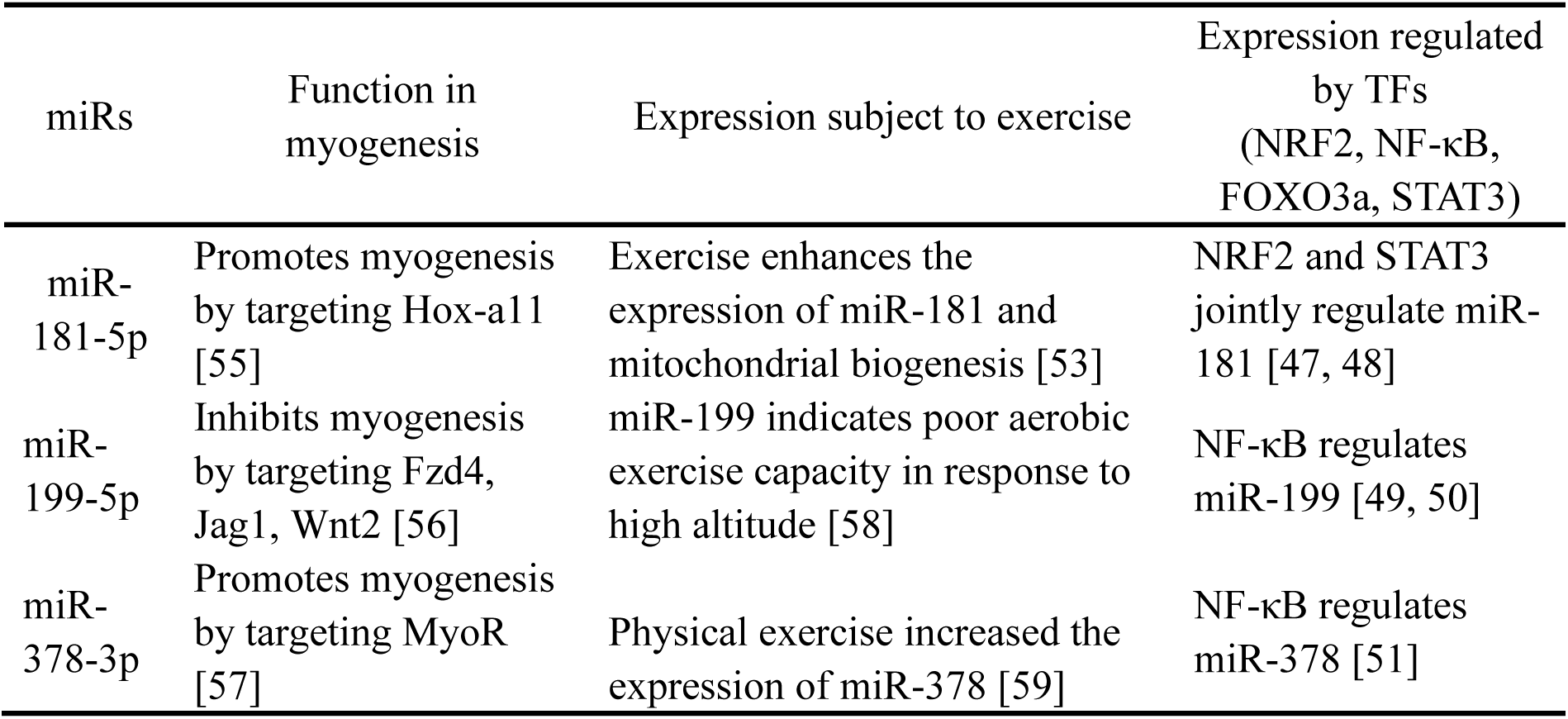
Selected miRs identified as having a role in exercise.

| miRs | Function in myogenesis | Expression subject to exercise | Expression regulated by TFs (NRF2, NF-κB, FOXO3a, STAT3) |
| --- | --- | --- | --- |
| miR-181-5p | Promotes myogenesis by targeting Hox-a11 [55] | Exercise enhances the expression of miR-181 and mitochondrial biogenesis [53] | NRF2 and STAT3 jointly regulate miR-181 [47, 48] |
| miR-199-5p | Inhibits myogenesis by targeting Fzd4, Jag1, Wnt2 [56] | miR-199 indicates poor aerobic exercise capacity in response to high altitude [58] | NF-κB regulates miR-199 [49, 50] |
| miR-378-3p | Promotes myogenesis by targeting MyoR [57] | Physical exercise increased the expression of miR-378 [59] | NF-κB regulates miR-378 [51] |

To evaluate the redox sensitivity of specific miRs in skeletal muscle, myoblasts were exposed to 25 µM H₂O₂ for 10 minutes and allowed to recover in fresh media for 24h. This concentration and duration have been previously shown to increase mitochondrial content and promote myogenesis [27]. qPCR analysis revealed a significant upregulation of mmu-miR-181a-5p, mmu-miR-199a-5p and mmu-miR-378a-3p at 24 hours post-exposure to acute H₂O₂ (Fig. 5a). Given the limited understanding of the roles of mmu-miR-181a-5p, mmu-miR-199a-5p and mmu-miR-378a-3p in regulating mitochondrial capacity and myogenesis in skeletal muscle, this study further explored their function by treating cells with miRs mimics (miR181, miR199, miR378) and antagomirs (AM181, AM199, AM378) prior to H₂O₂ exposure. Specifically, the miR181, miR199 and miR378 groups significantly increased, while AM181, AM199 and AM378 groups significantly decreased the levels of the respective miRs (Fig. 5a-c). The qPCR results consistently showed an increase in the expression of these miRs following H₂O₂ treatment (Fig. 5a-c). Additionally, following H₂O₂ treatment, mmu-miR-199a-5p and mmu-miR-378a-3p levels decreased in the miR199 and miR378 treated groups (Fig. 5b-c).

**Figure 5.**
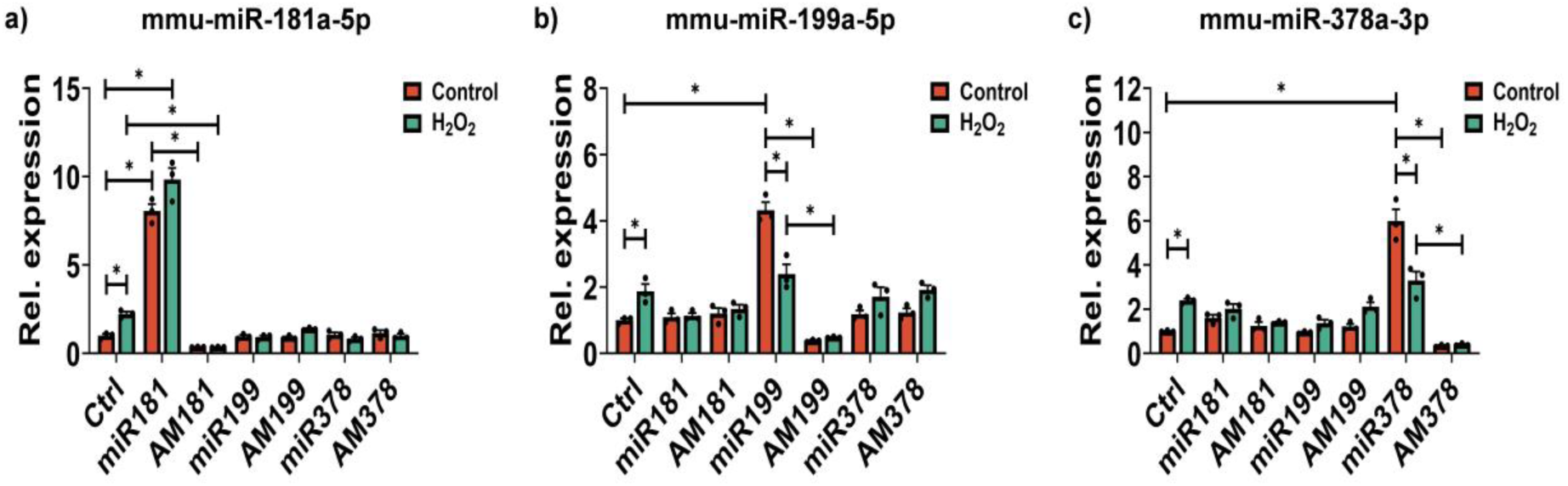
mmu-miR-181a-5p, mmu-miR-199a-5p and mmu-miR-378a-3p are up-regulated following an acute physiological concentration of H_2_O_2_ in skeletal muscle. C2C12 cells were transfected with mmu-miR-181a-5p, mmu-miR-199a-5p, mmu-miR-378a-3p mimics and antagomirs following physiological levels of H_2_O_2_ treatment (a-c). Graphs are the normalised relative means ± SEM and all experiments were performed with n = 3 and *p*-value of < 0.05 was considered as statistically significant *(p < 0.05). *p* values (a: in control group: ctrl vs miR181 < 0.0001, miR181 vs AM181 < 0.0001; in H_2_O_2_ group: ctrl vs AM181 = 0.0001; Control: ctrl vs H_2_O_2_: ctrl = 0.0264, Control: ctrl vs H_2_O_2_: miR181 < 0.0001. b: in control group: ctrl vs miR199 < 0.0001, miR199 vs AM199 < 0.0001; in H_2_O_2_ group: miR199 vs AM199 < 0.0001; Control: ctrl vs H_2_O_2_: ctrl = 0.0423, Control: ctrl vs H_2_O_2_: miR199 = 0.0001, Control: miR199 vs H_2_O_2_: miR199 < 0.0001. c: in control group: ctrl vs miR378 < 0.0001, miR378 vs AM378 < 0.0001; in H_2_O_2_ group: miR378 vs AM378 < 0.0001; Control: ctrl vs H_2_O_2_: ctrl = 0.0047, Control: miR378 vs H_2_O_2_: miR378 < 0.0001)

### 3.6 miR181, AM199 and miR378 increased mitochondrial content and promoted myogenesis

The potential regulatory roles of mmu-miR-181a-5p, mmu-miR-199a-5p and mmu-miR-378a-3p on mitochondrial content biogenesis, ROS and myogenesis were explored. In the control group, Western blot analysis in the control group revealed that the miR181, miR378 and AM199 groups led to increased expression of key mitochondrial regulatory proteins, including TOM20, VDAC and SIRT1 (Fig. 6a-c). In the H_2_O_2_ treated group, elevated levels of TOM20, VDAC and SIRT1 were also observed. However, a notable reduction in these protein levels was observed in the miR378 group following H₂O₂ treatment. Previously it has been reported that mmu-miR-181a-5p and mmu-miR-378a-3p play a role in autophagy regulation in mammalian cells [54, 60]. In the control group, the miR181 group had decreased LC3B-II/I ratio, whereas the AM181 group exhibited an increased ratio (Fig. S4b). Moreover, the miR378 group exhibited an elevated LC3b II/I ratio, and increased ATG5 and ULK1, along with a decrease in p62 levels, these findings indicated the role of mmu-miR-181a-5p and mmu-miR-378a-3p in regulating autophagy [60, 61]. The results highlight that an acute physiological concentration of H₂O₂ and that mmu-miR-181a-5p and mmu-miR-378a-3p, promote autophagy in myoblasts.

**Figure 6.**
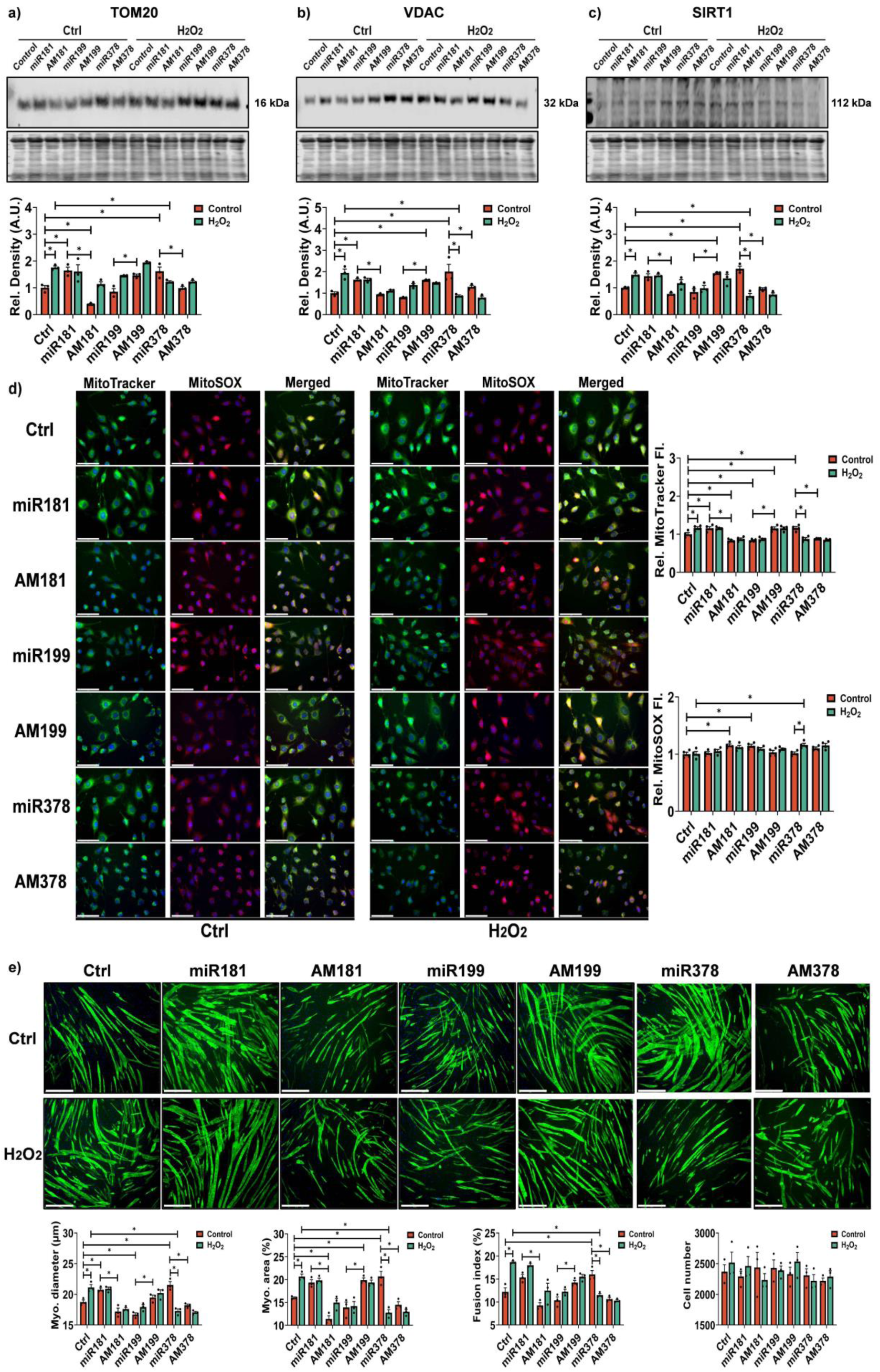
mmu-miR-181a-5p, mmu-miR-199a-5p and mmu-miR-378a-3p regulate mitochondrial biogenesis and myogenesis in skeletal muscle. Western blot indicating changes in the protein levels of mitochondria-associated proteins TOM20, VDAC and SIRT1 (a-c), autophagy-associated proteins p62, LC3b, ATG5 and ULK1 (d-g); MitoTracker and MitoSOX immunostaining showing changes of mitochondrial content and ROS levels (h); MF20 immunostaining confirming changes in myotube formation (i). Graphs are the normalised relative means ± SEM and all experiments were performed with n = 3-4 and *p*-value of < 0.05 was considered as statistically significant *(p < 0.05). *p* values (a: in control group: ctrl vs miR181 = 0.0063, ctrl vs AM181 = 0.0122, ctrl vs miR378 = 0.0103, miR181 vs AM181 < 0.0001, miR199 vs AM199 = 0.0121, miR378 vs AM378 = 0.0088; Control: ctrl vs H_2_O_2_: ctrl = 0.0009; in H_2_O_2_ group: ctrl vs miR378 = 0.0442. b: in control group: ctrl vs miR181 = 0.025, ctrl vs AM199 = 0.0335, ctrl vs miR378 < 0.0001, miR181 vs AM181 = 0.0094, miR199 vs AM199 = 0.0013, miR378 vs AM378 = 0.0066; in H_2_O_2_ group: ctrl vs miR378 < 0.0001; Control: ctrl vs H_2_O_2_: ctrl = 0.0001; Control: miR378 vs H_2_O_2_: miR378 < 0.0001. c: in control group: ctrl vs AM199 = 0.0082, ctrl vs miR378 = 0.0003, miR181 vs AM181 = 0.0008, miR199 vs AM199 = 0.0003, miR378 vs AM378 = 0.0001; Control: ctrl vs H_2_O_2_: ctrl = 0.0282, Control: miR378 vs H_2_O_2_: miR378 < 0.0001. d: MitoTracker: in control group: ctrl vs miR181 = 0.0346, ctrl vs AM181 = 0.0196, ctrl vs miR199 = 0.0269, ctrl vs AM199 = 0.0462, ctrl vs miR378 = 0.0229, miR181 vs AM181 < 0.0001, miR199 vs AM199 < 0.0001, miR378 vs AM378 < 0.0001; Control: ctrl vs H_2_O_2_: ctrl = 0.0248, Control: miR378 vs H_2_O_2_: miR378 < 0.0001. MitoSOX: in control group: ctrl vs AM181 = 0.024, ctrl vs miR199 = 0.0418, ctrl vs miR199 = 0.0418; Control: ctrl vs H_2_O_2_: miR378 = 0.0176; Control: miR378 vs H_2_O_2_: miR378 = 0.05. e: Myotube diameter: in control group: ctrl vs miR181 = 0.0401, ctrl vs miR199 = 0.0329, ctrl vs miR378 = 0.0011, miR181 vs AM181 < 0.0001, miR199 vs AM199 = 0.0011, miR378 vs AM378 < 0.0001; Control: ctrl vs H_2_O_2_: ctrl = 0.0077, Control: ctrl vs H_2_O_2_: miR378 < 0.0001, Control: miR378 vs H_2_O_2_: miR378 < 0.0001. Myotube area: in control group: ctrl vs AM181 = 0.0073, ctrl vs AM199 = 0.0462, ctrl vs miR378 = 0.0072; miR181 vs AM181 < 0.0001, miR199 vs AM199 = 0.0003, miR378 vs AM378 = 0.0002; Control: ctrl vs H_2_O_2_: ctrl = 0.0076; H_2_O_2_: ctrl vs H_2_O_2_: miR378 < 0.0001; Control: miR378 vs H_2_O_2_: miR378 < 0.0001. Fusion index: in control group: ctrl vs miR378 = 0.0302, miR181 vs AM181 < 0.0001, miR199 vs AM199 = 0.0249, miR378 vs AM378 = 0.0005; Control: ctrl vs H_2_O_2_: ctrl < 0.0001, H_2_O_2_: ctrl vs H_2_O_2_: miR378 < 0.0001, Control: miR378 vs H_2_O_2_: miR378 = 0.0049)

In order to determine the effects of mmu-miR-181a-5p, mmu-miR-199a-5p and mmu-miR-378a-3p on mitochondria and myogenesis, we performed MitoTracker green staining for mitochondrial content, MitoSOX red staining for mitochondrial ROS and MF20 staining to determine the effects on myogenesis following the acute physiological concentration of H_2_O_2_ treatment. In the control groups, the miR181, AM199 and miR378 groups exhibited increased MitoTracker green staining (Fig. 6d), as well as enhanced myotube diameter, myotube area and fusion index (Fig. 6e). However, the AM181 and miR199 groups demonstrated increased MitoSOX fluorescence intensity, along with decreased myotube diameter and myotube area. In the H_2_O_2_-treated groups, myoblasts exhibited heightened mitochondrial content and myogenesis, consistent with the observed mitochondrial regulatory protein levels (Fig. 6a-c). However, in the miR378 group treated with H_2_O_2_, a decrease in MitoTracker green intensity, an increase in MitoSOX intensity and impaired myogenesis were observed. Collectively, these findings suggest that mmu-miR-181a-5p, mmu-miR-199a-5p and mmu-miR-378a-3p play crucial roles in regulating mitochondrial content, autophagy and myogenesis.

### 3.7 Exercise-related cel-miR-77-5p and cel-miR-249-3p in *C. elegans* display conserved orthologous targets with exercise-related mmu-miR-181a-5p and mmu-miR-378a-3p

Although the exercise-related miRs identified in *C. elegans* and murine myoblasts are not conserved, we hypothesised that these miRs may share similar targets involved in regulating key mechanisms of muscle function. In a previous study, we performed a proteomics analysis following exercise in the N2 wild type strain, identifying 3,590 proteins, with 66 upregulated and 67 downregulated proteins compared to non-exercised controls [27]. A comparative analysis between the predicted targets of cel-miR-77-5p and cel-miR-249-3p and the proteomics data was performed. This analysis identified 51 predicted targets for cel-miR-77-5p, including 21 upregulated and 25 downregulated proteins, and 19 predicted targets for cel-miR-249-3p, encompassing 8 downregulated and 11 upregulated proteins (Fig. S5a). Within the predicted targets of cel-miR-77-5p, ten proteins associated with muscle contraction were identified: NCX-2, SCA-1, IPP-5, KIN-1, SCPL-4, LMP-1, CPZ-2, UNC-112, GRD-5 and T09B4.5. For cel-miR-249-3p, five proteins related to muscle contraction were highlighted: LAF-1, KSR-2, UNC-54, MAX-2 and PUF-9. Further analysis using TargetScan (https://www.targetscan.org) identified the top four potential targets of cel-miR-77-5p (*ncx-2*, *sca-1*, *ipp-5*, *kin-1*) and two potential targets of cel-miR-249-3p (*laf-1*, *ksr-2*) (Table 2). Subsequent qPCR analysis in the *C. elegans* N2 wild type strain confirmed that following exercise there was an upregulation of genes involved in intracellular Ca^2+^ regulation, *ipp-5*, *ncx-2* and *sca-1* (Fig. 7a,b,d). In the *mir-77* mutant strain, there was basally elevated levels of *ipp-5*, *ncx-2* and *kin-1* compared to N2 (Fig. 7a-c) suggesting that they are potential targets of cel-miR-77. For cel-miR-249-3p, qPCR analysis demonstrated elevated expression levels of *laf-1* and *ksr-2* in the *mir-249* mutant strain compared to N2 (Fig. 7e-f). These results suggest that *ncx-2*, *kin-1* and *ipp-5* are potential targets of cel-miR-77-5p, whereas *laf-1* and *ksr-2* are potential targets of cel-miR-249-3p.

**Figure 7.**
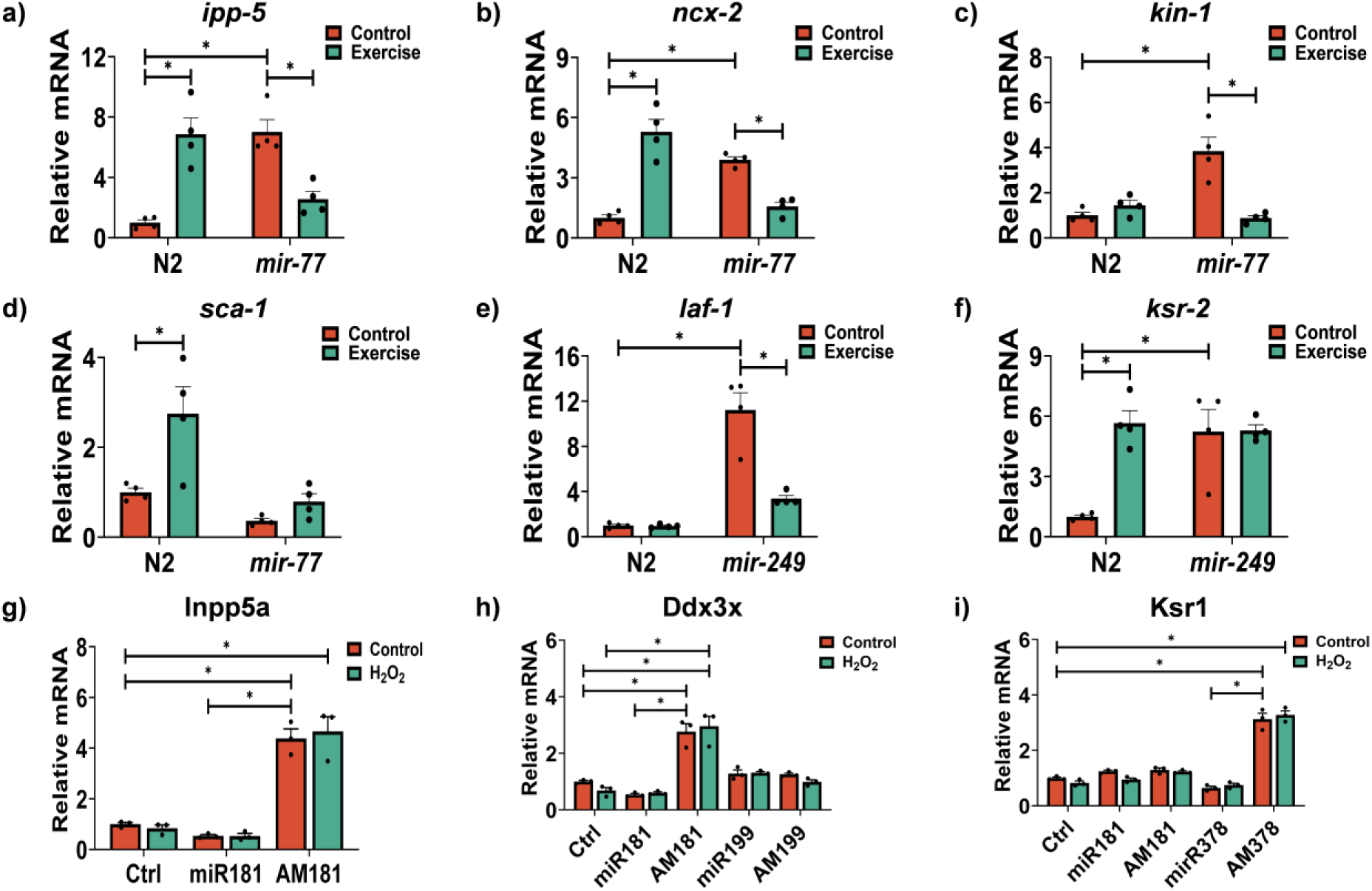
Target genes regulated by the exercise-related miRs in *C. elegans* and mammalian cells. Expression levels of predicted targets in *C. elegans* including *ipp-5* (a), *ncx-2* (b), *kin-1* (c), *sca-1* (d), *laf-1* (e), *ksr-2* (f) and predicted targets in mice including Inpp5a (g), Ddx3x (h), Ksr1 (i). Graphs are the normalised relative means ± SEM and all experiments were performed with n = 4 and *p*-value of < 0.05 was considered as statistically significant *(p < 0.05). *p* values (a: N2 vs *mir-77* = 0.0004; N2: Control vs Exercise = 0.0005; *mir-77*: Control vs Exercise = 0.0046. b: N2 vs *mir-77* = 0.0003; N2: Control vs Exercise < 0.0001; *mir-77*: Control vs Exercise = 0.0023. c: N2 vs *mir-77* = 0.0003; *mir-77*: Control vs Exercise = 0.0002. d: N2: Control vs Exercise = 0.01. e: N2 vs *mir-249* < 0.0001; *mir-249*: Control vs Exercise < 0.0001. f: N2 vs *mir-249* = 0.0028; N2: Control vs Exercise = 0.0014. g: in control group: ctrl vs AM181 < 0.0001, miR181 vs AM181 < 0.0001; Control: ctrl vs H_2_O_2_: AM181 < 0.0001. h: in control group: ctrl vs AM181 < 0.0001, miR181 vs AM181 < 0.0001; in H_2_O_2_ group: ctrl vs AM181 < 0.0001; Control: ctrl vs H_2_O_2_: AM181 < 0.0001. i: in control group: ctrl vs AM378 < 0.0001, miR378 vs AM378 < 0.0001; Control: ctrl vs H_2_O_2_: AM378 < 0.0001)

**Table 2.**
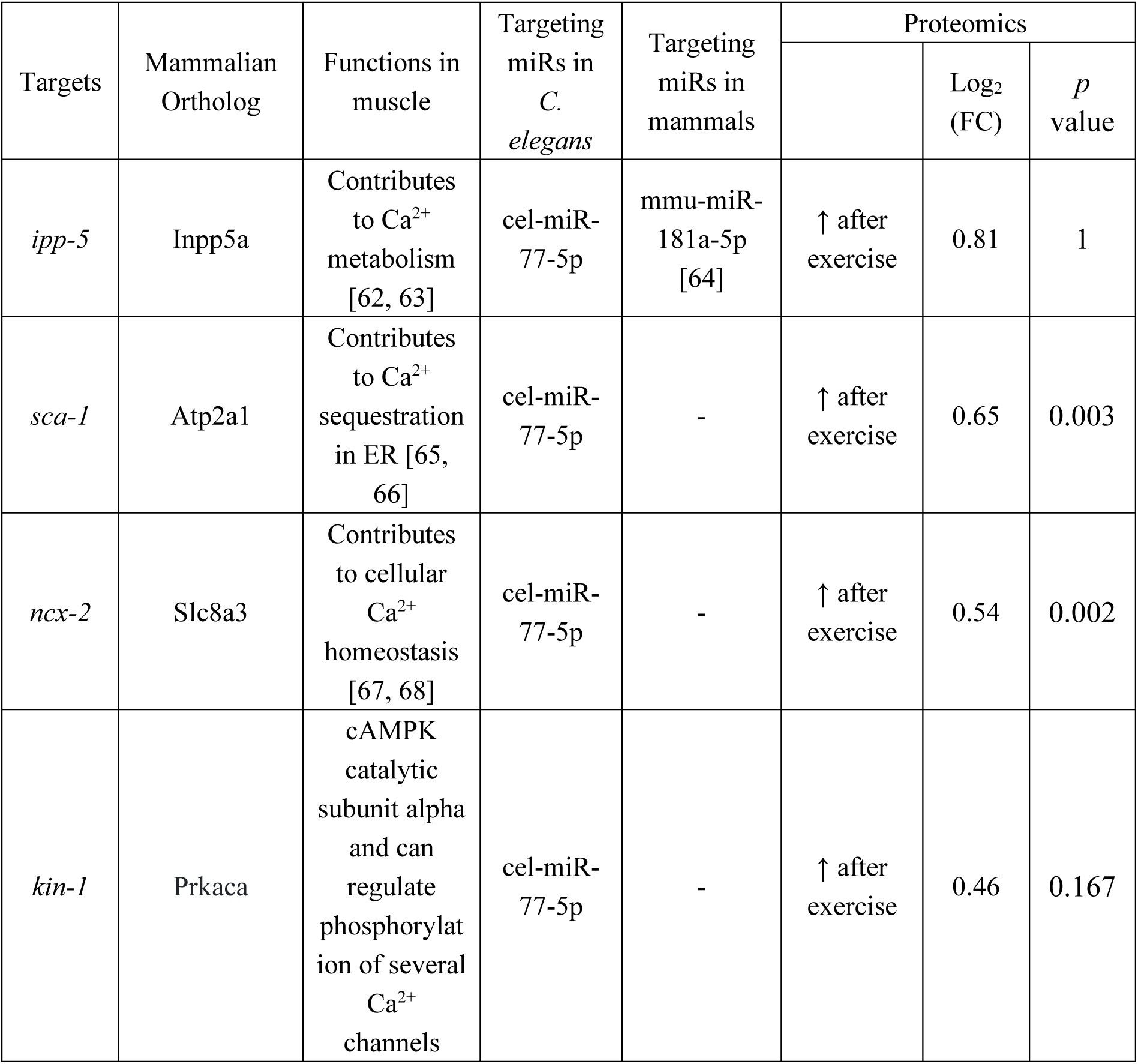
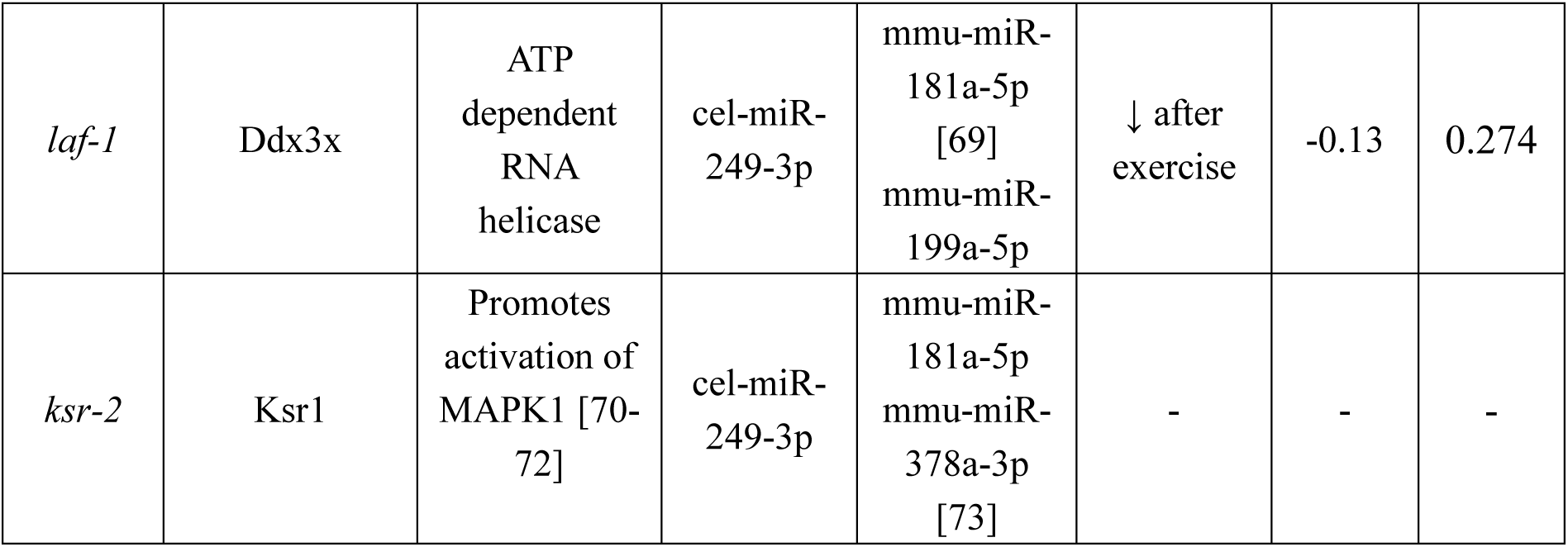
Summary of interested targets with known or potential role in muscle and their binding miRs.

While certain miRs are conserved across species [74], many exhibit species-specific sequences or significant divergence. Our data indicated that mmu-miR-181a-5p, mmu-miR-199a-5p and mmu-miR-378a-3p in mice share analogous physiological roles with cel-miR-77-5p and cel-miR-249-3p in *C. elegans*, particularly in regulating Ca^2+^ homeostasis and metabolic pathways. To explore conserved molecular targets, we utilised WORMHOLE (https://wormhole.jax.org/) [75], to identify orthologous genes targeted by cel-miR-77-5p and cel-miR-249-3p in *C. elegans* and mmu-miR-181a-5p, mmu-miR-199a-5p and mmu-miR-378a-3p in mice, thus revealing shared targets between the two species (Fig. S5b-c). Our analysis identified that cel-miR-77-5p shares 12 common targets with mmu-miR-181a-5p, 2 with mmu-miR-199a-5p and 2 with mmu-miR-378a-3p. Similarly, cel-miR-249-3p exhibits 16 shared targets with mmu-miR-181a-5p, 9 with mmu-miR-199a-5p and 2 with mmu-miR-378a-3p. Interestingly, Inpp5a (Inositol Polyphosphate-5-Phosphatase A) an ortholog of *ipp-5*, emerged as a shared target between cel-miR-77-5p and mmu-miR-181a-5p. Ddx3x (DEAD-box helicase 3 X-linked), an ortholog of *laf-1*, was identified as a common target of cel-miR-249-3p and both mmu-miR-181a-5p and mmu-miR-199a-5p, while Ksr1 (Kinase Suppressor of Ras), an ortholog of *ksr-2*, was a shared target of cel-miR-249-3p between mmu-miR-181a-5p and mmu-miR-378a-3p. To validate these putative shared targets, qPCR confirmed an increased expression of Inpp5a in the AM181 group, both with and without H₂O₂ treatment (Fig. 7g). Additionally, Ddx3x expression was upregulated following mmu-miR-181a-5p inhibition, but no changes were observed with mmu-miR-199a-5p manipulation (Fig. 7h). Moreover, Ksr1 levels were elevated under mmu-miR-378a-3p inhibition, with no significant correlation observed with mmu-miR-181a-5p (Fig. 7i). These findings suggest a conserved regulatory mechanism where cel-miR-77-5p and mmu-miR-181a-5p target *ipp-5*/Inpp5a, cel-miR-249-3p and mmu-miR-181a-5p share *laf-1*/Ddx3x, cel-miR-249-3p and mmu-miR-378a-3p share *ksr-2*/Ksr1. This comparative approach sheds light on evolutionary conserved mechanisms underlying physiological adaptations to exercise, revealing conserved molecular targets regulated by exercise-related miRs across species.

### 3.8 AM181 pretreatment decreases Ca^2+^ release in myotubes

Ca^2+^ homeostasis and regulation is key during contractile activity and for the adaptive response to exercise. INPP5a is a negative regulator of IP_3_ resulting in decreased Ca^2+^ release from the ER. The Na^+^/Ca^2+^ pump NCX-3, can also regulate intracellular Ca^2+^ levels and prevent Ca^2+^ overload during conditions of ischaemia and hypoxia. cel-miR-77-5p is predicted to target both *ipp-5* the *C. elegans* ortholog of Inpp5a and *ncx-2* the ortholog of the Na^+^/Ca^2+^ exchanger. Following exercise, levels of *ipp-5* and *ncx-2* increased and the *mir-77* mutant strain was characterised by basally elevated levels of *ipp-5* and *ncx-2* compared to the N2 strain (Fig. 7). As cel-miR-77-5p levels decreased following exercise (Fig. 1a) and there was a corresponding increase in *ipp-5* levels, indicating *ipp-5* as a target of cel-miR-77-5p (Fig. 7a). Inpp5a is also a predicted target of mmu-miR-181a-5p. In the C2C12 myoblasts, miR181 overexpression led to decreased levels of Inpp5a while AM181 treatment resulted in increased levels of Inpp5a (Fig. 7g). To determine the physiological consequences of altered Inpp5a levels, intracellular Ca^2+^ imaging was performed on myotubes that were pretreated with miR181 or AM181. ATP acting via G-protein coupled P2Y2R, results in a rapid transient spike in intracellular Ca^2+^ which was attenuated by suramin, an inhibitor of P2Y2R. Cells pre-treated with AM181 demonstrated a significantly decreased Ca^2+^ release profile in response to ATP, while miR181treatment resulted in a non-significant increase in Ca^2+^ release (Fig. 8a-b). Pre-incubation with suramin, resulted in an anticipated ∼50% decrease in Ca^2+^ transients and there was no effect of either miR181 or AM181 pretreatments, indicating that that changes observed were likely due to IP_3_ mediated Ca^2+^ signalling (Fig. 8c-d). We hypothesise that in the absence of suramin, AM181 resulted in increased levels of Inpp5a, which decreased IP_3_ that was observed as a reduction in Ca^2+^ transients. These results demonstrate that exercise related miRs, although not conserved, target physiologically relevant pathways associated with dynamic Ca^2+^ homeostasis and the adaptive response to exercise.

**Figure 8.**
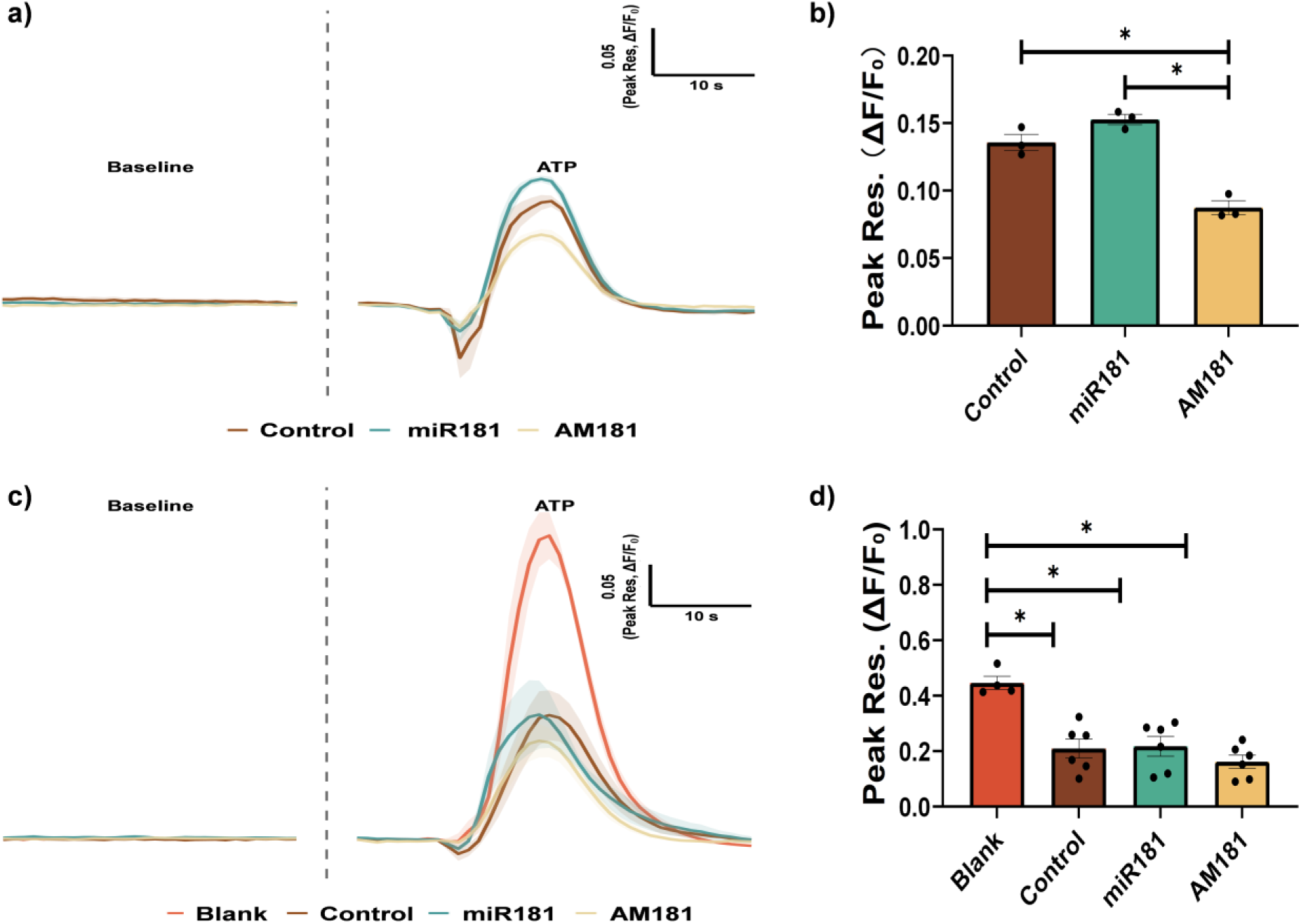
AM181 decreases Ca^2+^ release in myotubes. Ca^2+^ imaging performed without (a-b) or with Suramin (c-d). Graphs are the normalised relative means ± SEM and all experiments were performed with n = 3-6 and *p*-value of < 0.05 was considered as statistically significant *(*p* < 0.05). p values (b: Control vs AM181 = 0.0012; miR181 vs AM181 = 0.0002. d: Blank vs Control = 0.0005; Blank vs miR181 = 0.0007; Blank vs AM181 < 0.0001)

Small RNA sequencing revealed exercise-induced changes in miRs, increased levels of cel-miR-57-5p and cel-miR-249-3p but decreased levels of cel-miR-72-3p and cel-miR-77-5p. *mir-57* and *mir-249* mutant strains exhibited enhanced fertility, survival and lifespan, while *mir-72* and *mir-77* mutant strains demonstrated diminished fertility, survival and lifespan. Moreover, both *miR-77* and *mir-249* mutant strains exhibited reduced fitness and impaired mitochondrial morphology following exercise. While the seed sequences of the exercise responsive miRs identified in *C. elegans* are not conserved in mammals, literature-based identification of exercise-related mammalian miRs, such as mmu-miR-181a-5p, mmu-miR-199a-5p and mmu-miR-378a-3p were validated following acute H₂O₂ treatment of myoblasts. Increased levels of mmu-miR-181a-5p and mmu-miR-378a-3p enhanced mitochondrial content and myogenesis. Comparative analysis of predicted targets identified shared pathways, particularly in regulating Ca²⁺ homeostasis, with both mmu-miR-181a-5p and cel-miR-77-5p targeting Inpp5a/*ipp-5* to modulate Ca²⁺ metabolism. Ca^2+^ imaging in myotubes demonstrated decreased Ca²⁺ flux following AM181 treatment that was accompanied by increased Inpp5a levels, potentially decreasing IP_3_ mediated Ca^2+^ release. Additionally, mmu-miR-378a-3p and cel-miR-249-3p targeted Ksr1/*ksr-2*, involved in activating the MAPK pathway to regulate mitochondrial dynamics. These findings highlight conserved regulatory mechanisms linking miRs to Ca²⁺ homeostasis and mitochondrial function. Together, the data indicates common pathways respond to exercise in *C. elegans* and mammalian cells that are regulated by divergent miRs.

## 4 Discussion

Regular exercise helps maintain muscle mass and ameliorate age-related muscle atrophy. Exercise enhances mitochondrial biogenesis and the adaptive response to exercise results in the activation of redox sensitive transcription factors influencing gene expression [76]. However, the intricate mechanisms underlying the adaptive response to an acute physiological stress generated by exercise remain incompletely understood. Previous research has demonstrated that exercise enhances mitochondrial function, stress resistance and longevity in *C. elegans* [25, 27]. miRs have been identified as central regulators of post-transcriptional control in muscle proliferation, myogenesis and homeostasis [54, 57, 77]. In this study, we identified common pathways regulated by exercise related miRs. In *C. elegans*, a 5-day chronic exercise regimen enhanced mitochondrial content, survival, lifespan and fitness [27]. Following exercise, cel-miR-57-5p and cel-miR-249-3p were upregulated, while cel-miR-72-3p and cel-miR-77-5p were downregulated. Phenotypic analyses of *mir-57* and *mir-249* mutant strains revealed reduced fertility but increased survival and extended lifespan. Conversely, *mir-72* and *mir-77* mutants exhibited heightened fertility, decreased survival and shortened lifespan. In myoblasts, mmu-miR-181a-5p and mmu-miR-378a-3p enhanced mitochondrial capacity and myogenesis, whereas mmu-miR-199a-5p overexpression resulted in disrupted mitochondria and impaired myogenesis. Computational analyses identified that both mmu-miR-181a-5p and cel-miR-77-5p target Inpp5a/*ipp-5*, involved in regulating Ca^2+^ metabolism [62]. Furthermore, mmu-miR-378a-3p and cel-miR-249-3p were found to target Ksr1/*ksr-2*, activating the MAPK pathway that can regulate mitochondrial biogenesis [70].

The hormesis effect of exercise-induced ROS has been widely documented [78, 79]. A short bolus of H_2_O_2_ to myoblasts induced beneficial adaptive responses, including mitochondrial biogenesis and myogenesis [27]. Conversely, chronically elevated ROS levels can be deleterious, causing oxidative damage and impairing mitochondrial function [78]. Previously it was observed that acute H_2_O_2_ treatment induces the nuclear localisation of redox-sensitive transcription factors such as NRF2, NFκB and FOXO3, all of which have been associated with mitochondrial biogenesis in response to exercise [27]. Additionally, many of the exercise specific miRs contain binding sites for these redox regulated transcription factors (refer to Table 1 for details). Previous studies have established the regulatory role of miR-181 in mitochondrial homeostasis [80] and identified mmu-miR-181a-5p as a regulator of mitochondrial dynamics, targeting p62/SQSTM1, Protein DJ-1 and Parkin in skeletal muscle [54]. Additionally, miR-199-5p has been linked to smooth muscle hypertrophy [81], with elevated levels in human dystrophic muscle and impaired myofibers in zebrafish [56]. Recent research has also associated elevated miR-199-5p with cardiovascular dysfunction and reduced exercise endurance at high altitudes [58]. These findings suggest that miR-199-5p may serve as a negative regulator of muscle development, consistent with our findings. miR-378 has been reported to regulate glucose tolerance [82], mitochondrial homeostasis [83] and muscle development [84]. Furthermore, miR-378 has been identified as a promoter of autophagy in skeletal muscle [60], supporting the data obtained here. Unexpectedly, following H_2_O_2_ treatment, the miR378 group demonstrated compromised mitochondrial capacity, diminished myogenesis and heightened mitochondrial ROS levels, indicating a shift from the anticipated positive effects of mmu-miR-378a-3p to deleterious outcomes in myoblasts. A similar phenomenon was observed in *C. elegans*, particularly with cel-miR-249-3p, where its absence resulted in extended lifespan and survival, but increased mitochondrial fragmentation, decreased stress resistance and longevity following chronic exercise.

Small RNA sequencing revealed miRs affected by exercise in N2 and *prdx-2* mutant strains. Exercise in the N2 strain led to increased levels of 5 miRs, whereas the *prdx-2* mutant had increased levels of 14 miRs. The small RNA sequencing indicated that cel-miR-77-5p exhibited the most pronounced decrease in both N2 and *prdx-2* mutants following exercise. To identify potential targets of cel-miR-77-5p, we compared its predicted targets via TargetScan with the upregulated proteins identified in our previous proteomics data [27]. A number of potential targets were identified including, *ncx-2*, the ortholog of solute carrier family 8 member A3 (Slc8a3), that plays a crucial role in cellular Ca^2+^ homeostasis in muscle [67, 68]; *sca-1*, ortholog of ATPase sarcoplasmic/endoplasmic reticulum Ca^2+^ transporting 1 (Atp2a1), involved in Ca^2+^ sequestration during excitation-contraction coupling [65, 66]; Additionally, *ipp-5*, the ortholog of Inpp5a, plays a significant role in muscle function by hydrolysing IP_3_ to regulate Ca^2+^ signalling [63, 85].

Inpp5a was previously identified as a predicted target of mmu-miR-181 in HeLa cells [64] and our results indicate that its ortholog in *C. elegans*, *ipp-5,* is a target of cel-miR-77-5p. Inpp5a encodes the inositol polyphosphate-5-phosphatase enzyme critical for Ca^2+^ signalling, membrane trafficking and cell proliferation in skeletal muscle [86–88]. Comparing the predicted targets of cel-miR-249-3p, with previous quantitative proteomics data [27], four predicted genes are associated with mitochondrial function in skeletal muscle. These include, *laf-1*, the ortholog of Ddx3x, an ATP dependent RNA helicase which is implicated in mitochondrial DNA depletion and DDX3X syndrome [89], *unc-54*, the ortholog of Myh7, a motor protein with ATPase activity essential for muscle contraction [90, 91], *max-2*, the ortholog of Pak1, required for skeletal muscle mitochondrial function [92]; and *puf-9*, the ortholog of Pum2, which regulates mitochondrial dynamics and mitophagy [93]. Among these, Ddx3x/*laf-1* emerges as a shared target of both mmu-miR-181 and mmu-miR-199 [69], our data confirmed Ddx3x as a target of mmu-miR-181a-5p in myoblasts. Additionally, Ksr1/*ksr-2* is a shared predicted target of both mmu-mmu-miR-181a-5p and mmu-miR-378, as identified via TargetScan. qPCR result corroborated Ksr1 as a target of mmu-miR-378a-3p, aligning with previous findings that identified Ksr1 as a target of miR-378 in rat cardiomyocytes [73]. Ksr1 is a scaffold protein integral to the regulation of the MAPK signalling pathway, which plays a pivotal role in the adaptive response to exercise, encompassing critical aspects such as mitochondrial capacity and muscle function [71, 72].

16 miRs have been suggested as muscle-specific in the body wall muscle of *C. elegans* [94]. These include miR-241, miR-230, miR-77, miR-5551, miR-34, miR-228, miR-let-7, miR-80, miR-250, miR-239a, miR-239b, miR-392, miR-270, miR-4937, miR-793 and miR-799 [94]. Following chronic exercise, three of these miRs (cel-miR-77-5p, miR-80 and miR-250) were down-regulated in both N2 and *prdx-2* mutant strains. Interestingly, in the absence of PRDX-2, there was increased level of miR-34, a muscle-specific miR and an ortholog of vertebrate miR-34. miR-34 is recognised as a senescence-associated miR as its overexpression has been shown to induce cellular senescence [95], suppress muscle development [96] and impair insulin signalling pathway [97], mirroring similar phenotypes observed in the *prdx-2* mutant [27, 98, 99]. It is worth mentioning that three miRs (miR-37, miR-39 and miR-72) exhibited divergent changes in expression following exercise: decreased in the N2 wild type strain but increased in the *prdx-2* mutant strain. Specifically, miR-37-5p switched to miR-37-3p, miR-39-5p switched to miR-39-3p and miR-72-3p switched to miR-72-5p, although the overall levels of the passenger strand were still less than the levels of its guide strand. Arm switching is a remarkably dynamic process that can vary across different tissues, developmental stages or cellular conditions [100]. However, the detailed molecular mechanisms underlying arm switching are not fully understood [101]. In the context of our experimental findings, two potential factors may contribute to the observed phenomenon. First, oxidative stress induced by exercise and the absence of PRDX-2 could contribute, as oxidative stress is known to regulate RNA processing and the activity of key enzymes involved in miR assembly [102]. Second, our analysis of prior proteomics data suggests a possible role for PRDX-2 in miR biogenesis [27]. However, these hypotheses require further investigations to elucidate the precise molecular mechanisms underlying the observed arm switching.

Exercise can dramatically change mitochondrial morphology and it has been demonstrated in *C. elegans* to initially induce mitochondrial fragmentation followed by mitochondrial fusion after a recovery period, a cycle that becomes disrupted during ageing [25, 28]. Exercise can also modulate inter organelle communication between mitochondria and other organelles including lysosomes, lipid stores and the ER/SR [28, 103]. Inter-organelle crosstalk and the exchange of material and information in response to biological perturbations is crucial in order to respond to changes in cellular homeostasis [104]. Communication between the ER and mitochondria at contact sites are required for the transfer of Ca^2+^, lipids and metabolites [105]. MERCS are dynamic structures that can respond to changes in the cellular homeostasis and help determine sites of mitochondrial fission and fusion ultimately regulating mitochondrial morphology but also Ca^2+^ and lipid homeostasis [104, 106, 107]. Ca^2+^ is essential for muscle contraction and mitochondrial metabolism, which are fundamental during exercise [108]. Regular exercise enhances skeletal muscle Ca^2+^ handling and improves running performance in mice [109]. Impaired Ca^2+^ homeostasis can lead to an imbalance in Ca^2+^ uptake, changes in mitochondrial membrane potential and shifts in fibre type, all of which collectively influence exercise capacity in mice [110]. Conversely, excessive Ca²⁺ accumulation is associated with induction of apoptotic signalling and muscle dystrophy [111]. Thus, maintaining Ca²⁺ balance is crucial for optimal muscle function and development. Our findings indicate that the exercise-induced reduction in cel-miR-77-5p expression likely represents an acute adaptive response to enhance the expression of its target genes (*ipp-5*, *ncx-2* and *kin-1*), improve Ca²⁺ metabolism and mitochondrial function. In contrast, the complete absence of cel-miR-77-5p in the *mir-77* mutant may disrupt the regulation of these targets or lead to their uncontrolled upregulation, resulting in compromised Ca²⁺ handling, apoptotic signalling and mitochondrial dysfunction. This dysregulation could contribute to decreased mitochondrial potential and reduced lifespan observed in the *mir-77* mutant following exercise. Therefore, while the downregulation of cel-miR-77-5p during exercise appears to facilitate adaptive responses, its absence in the *mir-77* mutant emphasises the critical role of cel-miR-77-5p in maintaining Ca²⁺ homeostasis and mitochondrial health over the long term.

MAPK represents a family of serine/threonine kinases that are critical components of evolutionarily conserved signalling pathways. In skeletal muscle, MAPK is indispensable for processes such as muscle differentiation, regeneration and adaptations to physiological stress [112]. Exercise-induced MAPK activation stimulates the mitochondrial regulator PGC-1α, thereby promoting mitochondrial biogenesis and facilitating muscle adaptation [113]. Additionally, Ddx3x has been implicated in mitochondrial DNA depletion in skeletal muscle, further underscoring the importance of these pathways in mitochondrial health [89]. Given that both *ksr-2* (the ortholog of Ksr1, which activates MAPK) and *laf-1* (the ortholog of Ddx3x) are targets of cel-miR-249-3p, the observed increase in cel-miR-249-3p following exercise appears paradoxical when considering the *mir-249* mutant exhibits increased mitochondrial membrane potential and extended lifespan. This contradiction suggests that the exercise-induced upregulation of cel-miR-249-3p might be an adaptive response aimed at fine-tuning mitochondrial capacity to maintain a balance in mitochondrial function. In contrast, the enhanced mitochondrial membrane potential and extended lifespan observed in the *mir-249* mutant indicate that the absence of cel-miR-249-3p leads to the upregulation or increased activity of its targets, including *ksr-2* and *laf-1*, potentially resulting in alterations in mitochondrial dynamics. Therefore, while cel-miR-249-3p may serve to modulate these processes under normal conditions, its absence in the *mir-249* mutant might disrupt this regulation, leading to more pronounced effects on mitochondrial capacity and longevity. In the future, muscle specific miR sequencing and Ca^2+^ imaging in *C. elegans*, would provide a more direct and comprehensive approach to of the underlying mechanisms involved in the adaptive response to exercise.

In summary, the results highlight the involvement of cel-miR-57-5p, cel-miR-72-3p, cel-miR-77-5p and cel-miR-249-3p in the regulation of fertility, survival and mitochondrial capacity in *C. elegans*. Similarly, mmu-miR-181a-5p, mmu-miR-199a-5p and mmu-miR-378a-3p play key regulatory roles on mitochondrial content and myogenesis in C2C12 myoblasts. Computational analysis and RNA validation identified Inpp5a/*ipp-5* as a shared target of mmu-miR-181a-5p and cel-miR-77-5p, associated with Ca²⁺ homeostasis in skeletal muscle. Additionally, Ksr1/*ksr-2* was confirmed as a shared target of mmu-miR-378a-3p and cel-miR-249-3p, which is linked to MAPK activation for the regulation of mitochondrial biogenesis. Collectively, this data underscores the critical roles of miRs in conserved regulatory mechanisms governing mitochondrial function across species.

## Acknowledgements

We would like to sincerely thank Gibran Pedraza for creating the volcano plot featured in the paper. QX (202006370047), JT (202306370005) and PL (202206370063) studentships are funded by the Chinese Scholarship Council (CSC), JCCM studentship is funded by the College of Nursing Medicine and Health Sciences, University of Galway.

## Author Contributions

Conceptualisation, BMcD, KW and QX; methodology, QX, JT, JC, PL, LQ, KW and BMcD; investigation, QX, JT, JC, PL and BMcD; resources, LQ, KW and BMcD; writing-original draft, QX and BMcD; writing-review and editing, QX, JT, PL, JC, LQ, KW and BMcD; supervision, LQ, KW and BMcD.

---

**Figure S1.**
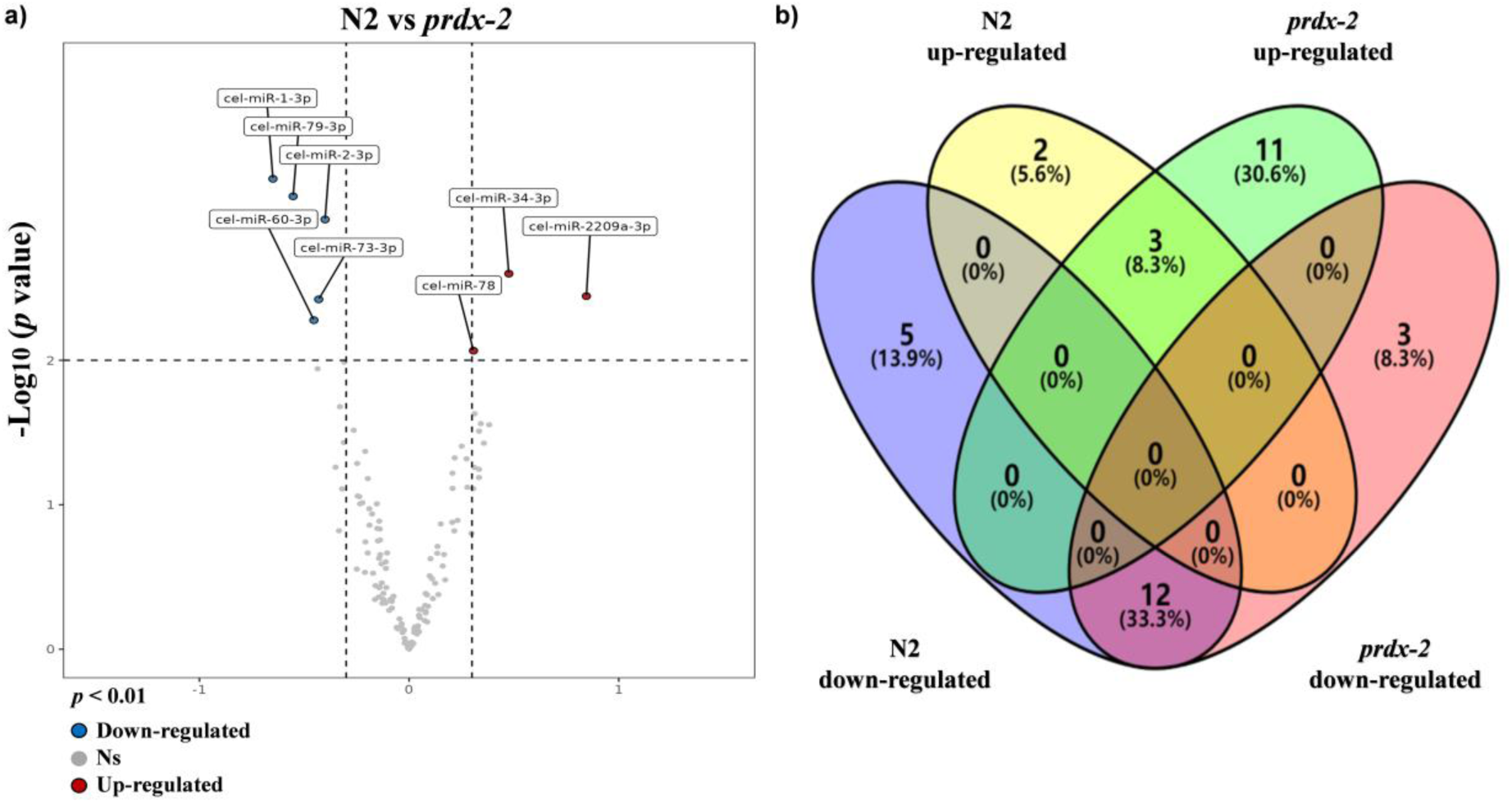
microRNA sequencing data following 5-days of swimming exercise. Non-exercised N2 vs non-exercised *prdx-2* mutant (FDR = 15.13%) (a); Venn diagram for the common altered miRs following exercise in N2 and *prdx-2* mutant (b).

**Figure S2.**
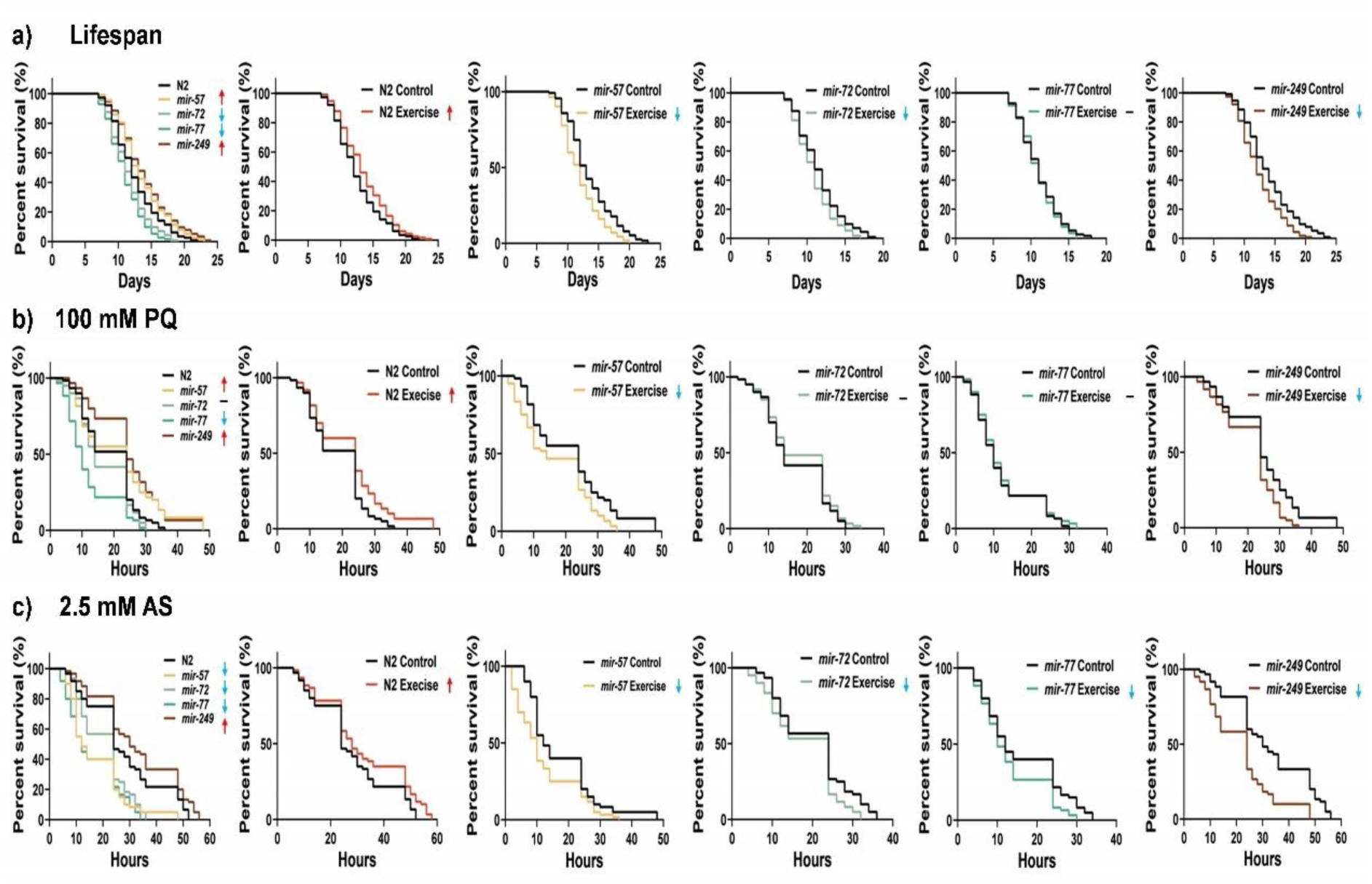
cel-miR-57-5p, cel-miR-72-3p, cel-miR-77-5p and cel-miR-249-3p regulate lifespan and resistance to oxidative stress in *C. elegans*. cel-miR-57-5p, cel-miR-72-3p, cel-miR-77-5p and cel-miR-249-3p regulate lifespan (a) and resistance to paraquat and arsenite (b-c). Graphs are the normalised relative means ± SEM and all experiments were performed with n=110-120 (lifespan assay), n=60 (paraquat and arsenite assay) and *p*-value of < 0.05 was considered as statistically significant *(*p* < 0.05). *p* values (a: N2 vs *mir-72* = 0.0233, N2 vs *mir-77* = 0.01, N2 vs *mir-249* = 0.0023. e: N2 vs *mir-57* = 0.0467, N2 vs *mir-72* = 0.0103, N2 vs *mir-77* = 0.0002, N2 vs *mir-249* = 0.0105; N2: Control vs Exercise = 0.0432; *mir-57*: Control vs Exercise = 0.0039; *mir-72*: Control vs Exercise = 0.0364; *mir-249*: Control vs Exercise = 0.0019. b: N2 vs *mir-57* = 0.0362, N2 vs *mir-77* < 0.0001, N2 vs *mir-249* = 0.0003; N2: Control vs Exercise = 0.0228; *mir-57*: Control vs Exercise = 0.0144; *mir-249*: Control vs Exercise = 0.0097. c: N2 vs *mir-57* < 0.0001, N2 vs *mir-72* = 0.0002, N2 vs *mir-77* < 0.0001, N2 vs *mir-249* = 0.0305; N2: Control vs Exercise = 0.0453; *mir-57*: Control vs Exercise = 0.0114; *mir-72*: Control vs Exercise = 0.0402; *mir-77*: Control vs Exercise = 0.0369; *mir-249*: Control vs Exercise < 0.0001)

**Figure S3.**
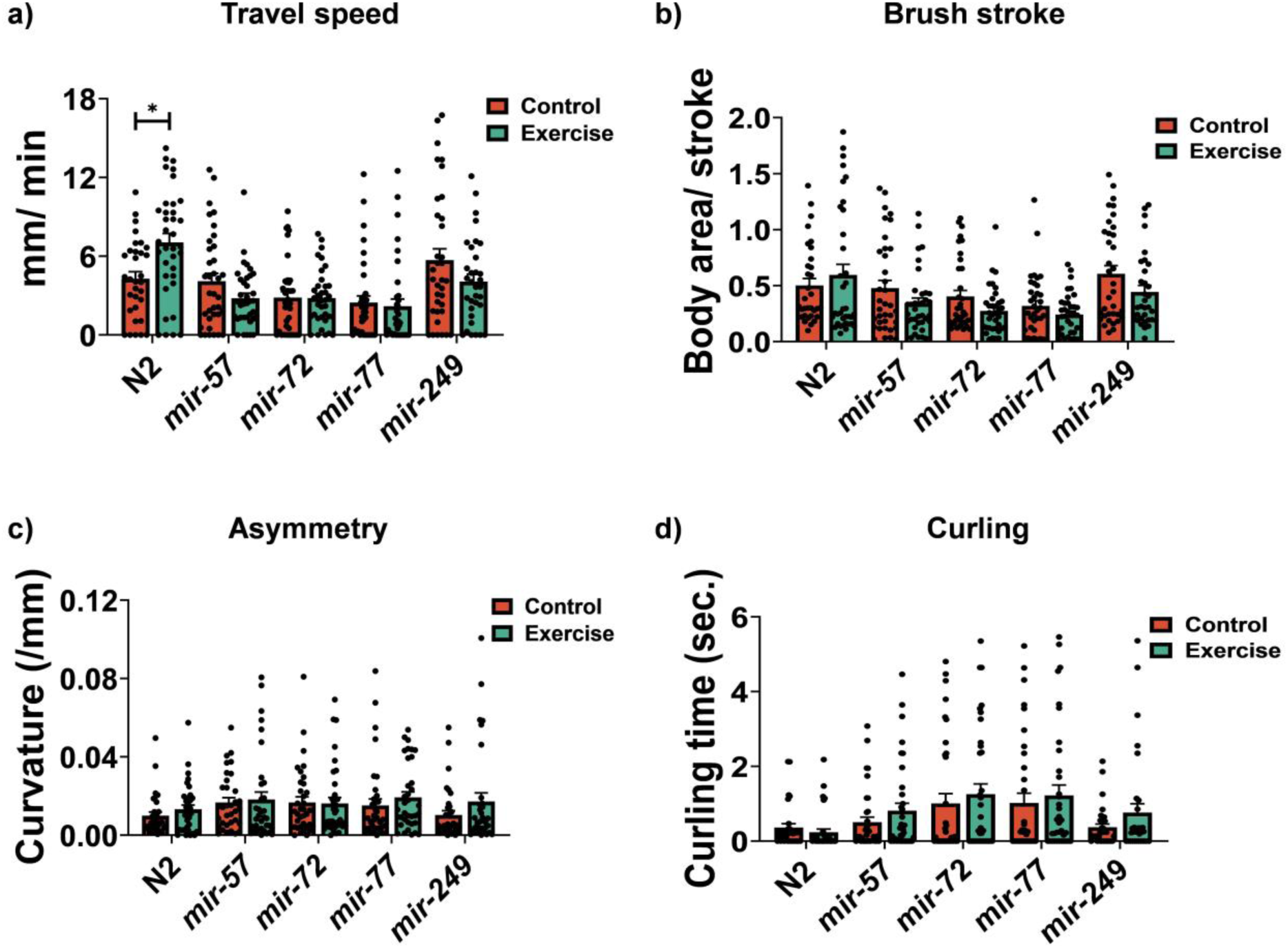
cel-miR-77-5p and cel-miR-249-3p are required for physical fitness following swimming exercise in *C. elegans*. Travel speed (a), brush stroke (b), asymmetry (c) and curling (d) following chronic swimming exercise. Graphs are the normalised relative means ± SEM and all experiments were performed with n = 30-40 and *p*-value of < 0.05 was considered as statistically significant *(p < 0.05). *p* values (a: N2: Control vs Exercise = 0.0224)

**Figure S4.**
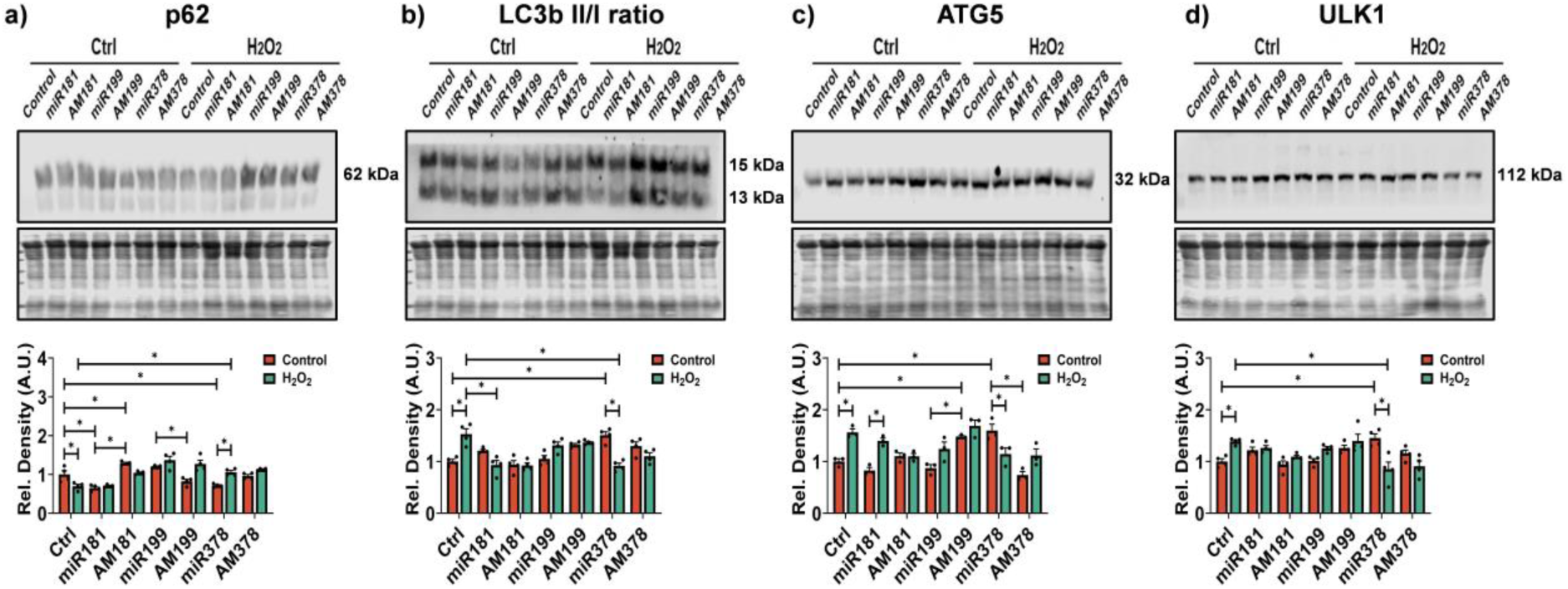
mmu-miR-181a-5p, mmu-miR-199a-5p and mmu-miR-378a-3p regulate autophagy levels in skeletal muscle. Western blot indicating changes in the protein levels of autophagy-associated proteins p62, LC3b, ATG5 and ULK1 (a-d). Graphs are the normalised relative means ± SEM and all experiments were performed with n = 3-4 and p-value of < 0.05 was considered as statistically significant *(p < 0.05). *p* values (a: in control group: ctrl vs miR181 = 0.0033, ctrl vs AM181 = 0.027, ctrl vs miR378 = 0.0317, miR181 vs AM181 < 0.0001, miR199 vs AM199 = 0.0011; in H_2_O_2_ group: ctrl vs miR378 = 0.0018; Control: ctrl vs H_2_O_2_: ctrl = 0.0163, Control: miR378 vs H_2_O_2_: miR378 = 0.0039. b: in control group: ctrl vs miR378 = 0.0001; Control: ctrl vs H_2_O_2_: ctrl < 0.0001; H_2_O_2_: ctrl vs H_2_O_2_: miR181 < 0.0001, H_2_O_2_: ctrl vs H_2_O_2_: miR378 < 0.0001, Control: miR378 vs H_2_O_2_:miR378 < 0.0001. c: in control group: ctrl vs AM199 = 0.0287, ctrl vs miR378 = 0.0025, miR199 vs AM199 = 0.0021, miR378 vs AM378 < 0.0001; Control: ctrl vs H_2_O_2_: ctrl = 0.0053; Control: miR378 vs H_2_O_2_: miR378 = 0.0445. d: in control group: ctrl vs miR378 = 0.0062; Control: ctrl vs H_2_O_2_: ctrl = 0.034; H_2_O_2_: ctrl vs H_2_O_2_: miR378 = 0.0005, Control: miR378 vs H_2_O_2_: miR378 < 0.0001)

**Figure S5.**
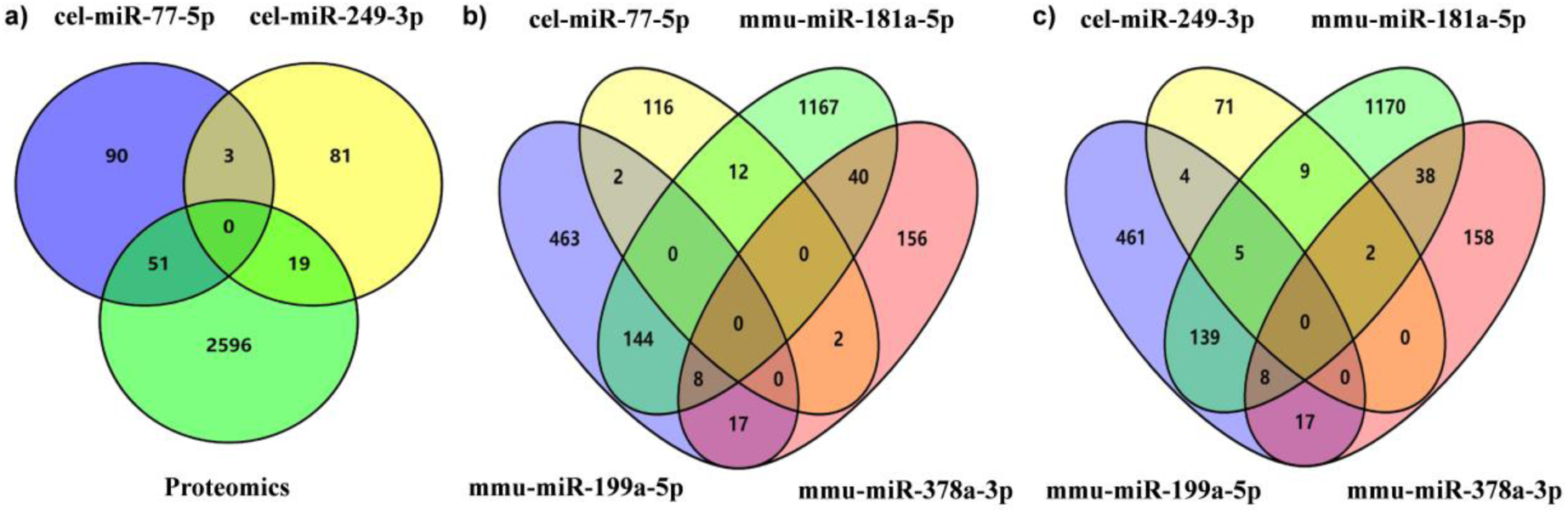
overlapping molecular targets that are potentially regulated by the exercise-related miRs in both species. Venn diagram of common targets among cel-miR-77-5p, cel-miR-249-3p and protein changes identified in the previous proteomics analysis (a), shared orthologous targets across cel-miR-77-5p, cel-miR-249-3p in *C. elegans* and mmu-miR-181a-5p, mmu-miR-199a-5p and mmu-miR-378a-3p in mouse (b,c).

**Table S1.** Significant changed miRs following exercise in N2 and *prdx-2* mutant.

| Conditions |  | microRNAs | log<sup>FC</sup> | *p* value |
| --- | --- | --- | --- | --- |
| N2<br>Control<br>vs<br>Exercise | up-regulated<br>miRs | cel-miR-249-3p | 0.7229 | 0.000387 |
|  |  | cel-miR-34-5p | 0.378 | 0.005743 |
|  |  | cel-miR-57-5p | 0.3583 | 0.007188 |
|  |  | cel-miR-52-5p | 0.3281 | 0.008151 |
|  |  | cel-miR-34-3p | 0.4654 | 0.00974 |
|  | down-regulated<br>miRs | cel-miR-77-5p | -2.233 | 1.001e-8 |
|  |  | cel-miR-39-5p | -1.273 | 4.21E-06 |
|  |  | cel-miR-41-5p | -1.005 | 8.43E-06 |
|  |  | cel-miR-58a-5p | -0.6944 | 0.000592 |
|  |  | cel-miR-80-5p | -0.7514 | 0.000635 |
|  |  | cel-miR-8202-5p | -0.6643 | 0.000654 |
|  |  | cel-miR-72-3p | -0.6125 | 0.001184 |
|  |  | cel-miR-229-3p | -0.4174 | 0.001996 |
|  |  | cel-miR-45-5p | -0.5903 | 0.002075 |
|  |  | cel-miR-37-5p | -0.8166 | 0.002777 |
|  |  | cel-miR-250-5p | -0.5631 | 0.002984 |
|  |  | cel-miR-238-5p | -0.6236 | 0.00306 |
|  |  | cel-miR-38-5p | -0.8243 | 0.003071 |
|  |  | cel-miR-36-5p | -0.6807 | 0.003205 |
|  |  | cel-miR-70-5p | -0.8919 | 0.007898 |
|  |  | cel-miR-71-3p | -0.4616 | 0.008123 |
|  |  | cel-miR-239a-3p | -0.484 | 0.008448 |
| *prdx2*<br>Control<br>vs<br>Exercise | up-regulated<br>miRs | cel-miR-57-5p | 0.5494 | 1.21E-05 |
|  |  | cel-miR-42-3p | 0.5399 | 1.68E-05 |
|  |  | cel-miR-37-3p | 0.5695 | 0.000113 |
|  |  | cel-miR-49-3p | 0.462 | 0.000167 |
|  |  | cel-miR-237-5p | 0.5375 | 0.000192 |
|  |  | cel-miR-52-5p | 0.3581 | 0.000256 |
|  |  | cel-miR-43-3p | 0.4896 | 0.000292 |
|  |  | cel-miR-231-3p | 0.3969 | 0.001446 |
|  |  | cel-miR-2-3p | 0.3703 | 0.001775 |
|  |  | cel-miR-74-3p | 0.6187 | 0.001871 |
|  |  | cel-miR-79-3p | 0.4641 | 0.005094 |
|  |  | cel-miR-249-3p | 0.6201 | 0.005397 |
|  |  | cel-miR-73-3p | 0.3986 | 0.005427 |
|  |  | cel-miR-76-3p | 0.3051 | 0.008625 |
|  | down-regulated<br>miRs | cel-miR-77-5p | -2.377 | 2.168e-10 |
|  |  | cel-miR-39-5p | -1.251 | 1.3E-05 |
|  |  | cel-miR-72-3p | -0.9696 | 2.39E-05 |
|  |  | cel-miR-37-5p | -1.027 | 0.000155 |
|  |  | cel-miR-58a-5p | -1.1 | 0.000164 |
|  |  | cel-miR-38-5p | -0.739 | 0.00019 |
|  |  | cel-miR-80-5p | -0.5562 | 0.000393 |
|  |  | cel-miR-41-5p | -0.7673 | 0.000683 |
|  |  | cel-miR-250-5p | -0.6796 | 0.000711 |
|  |  | cel-miR-71-3p | -0.6373 | 0.001036 |
|  |  | cel-miR-55-5p | -0.7181 | 0.001067 |
|  |  | cel-miR-78 | -0.427 | 0.002444 |
|  |  | cel-miR-36-5p | -0.5717 | 0.002499 |
|  |  | cel-miR-50-3p | -0.882 | 0.006353 |
|  |  | cel-miR-238-5p | -0.4793 | 0.009043 |
| N2 Control<br>vs<br>*prdx2* Control | up-regulated<br>miRs | cel-miR-34-3p | 0.4758 | 0.002516 |
|  |  | cel-miR-2209a-3p | 0.8458 | 0.003598 |
|  |  | cel-miR-78 | 0.3072 | 0.008586 |
|  | down-regulated<br>miRs | cel-miR-1-3p | -0.6494 | 0.000555 |
|  |  | cel-miR-79-3p | -0.5532 | 0.000733 |
|  |  | cel-miR-2-3p | -0.4007 | 0.001061 |
|  |  | cel-miR-73-3p | -0.4311 | 0.003788 |
|  |  | cel-miR-60-3p | -0.454 | 0.00528 |

**Table S2.** cel-miR-57-5p, cel-miR-72-3p, cel-miR-77-5p and cel-miR-249-3p regulate lifespan and resistance to oxidative stress in *C. elegans*.

| Figure | strain | Condition | # Deaths/<br># Total | Mean<br>lifespan | Log-rank test, *p*-value |
| --- | --- | --- | --- | --- | --- |
| 4d | N2 | Control | 113/120 | 12.76 ± 0.32 |  |
|  | N2 | Exercise | 115/120 | 13.75 ± 0.35 | compared to N2 C, *p* = 0.0432 |
|  | *mir-57* | Control | 113/120 | 13.73 ± 0.35 | compared to N2 C, *p* = 0.0467 |
|  | *mir-57* | Exercise | 112/120 | 12.36 ± 0.31 | compared to *mir-57* C, *p* = 0.0039 |
|  | *mir-72* | Control | 112/120 | 11.71 ± 0.27 | compared to N2 C, *p* = 0.0103 |
|  | *mir-72* | Exercise | 111/120 | 11.02 ± 0.23 | compared to *mir-72* C, *p* = 0.0364 |
|  | *mir-77* | Control | 112/120 | 11.2 ± 0.25 | compared to N2 C, *p* = 0.0002 |
|  | *mir-77* | Exercise | 114/120 | 10.95 ± 0.22 | compared to *mir-77* C, *p* = 0.3273 |
|  | *mir-249* | Control | 113/120 | 14.01 ± 0.36 | compared to N2 C, *p* = 0.0105 |
|  | *mir-249* | Exercise | 114/120 | 12.59 ± 0.3 | compared to *mir-249* C, *p* = 0.0019 |
| 4e | N2 | Control | 60/60 | 18.67 ± 1.08 |  |
|  | N2 | Exercise | 60/60 | 22.17 ± 1.42 | compared to N2 C, *p* = 0.0228 |
|  | *mir-57* | Control | 60/60 | 21.57 ± 1.61 | compared to N2 C, *p* = 0.0362 |
|  | *mir-57* | Exercise | 60/60 | 16.8 ± 1.38 | compared to *mir-57* C, *p* = 0.0144 |
|  | *mir-72* | Control | 60/60 | 16.73 ± 1.03 | compared to N2 C, *p* = 0.1772 |
|  | *mir-72* | Exercise | 60/60 | 17.87 ± 1.09 | compared to *mir-72* C, *p* = 0.3737 |
|  | *mir-77* | Control | 60/60 | 12.1 ± 0.99 | compared to N2 C, *p* < 0.0001 |
|  | *mir-77* | Exercise | 60/60 | 12.6 ± 1.02 | compared to *mir-77* C, *p* = 0.5456 |
|  | *mir-249* | Control | 60/60 | 24.77 ± 1.35 | compared to N2 C, *p* = 0.0003 |
|  | *mir-249* | Exercise | 60/60 | 21.07 ± 1.1 | compared to *mir-249* C, *p* = 0.0097 |
| 4f | N2 | Control | 60/60 | 28.13 ± 1.8 |  |
|  | N2 | Exercise | 60/60 | 32.23 ± 2.06 | compared to N2 C, *p* = 0.0453 |
|  | *mir-57* | Control | 60/60 | 17.4 ± 1.39 | compared to N2 C, *p* < 0.0001 |
|  | *mir-57* | Exercise | 60/60 | 12.17 ± 1.23 | compared to *mir-57* C, *p* = 0.0114 |
|  | *mir-72* | Control | 60/60 | 20.47 ± 1.15 | compared to N2 C, *p* = 0.0002 |
|  | *mir-72* | Exercise | 60/60 | 18.17 ± 1.09 | compared to *mir-72* C, *p* = 0.0402 |
|  | *mir-77* | Control | 60/60 | 16.13 ± 1.26 | compared to N2 C, *p* < 0.0001 |
|  | *mir-77* | Exercise | 60/60 | 13.33 ± 1.02 | compared to *mir-77* C, *p* = 0.0369 |
|  | *mir-249* | Control | 60/60 | 32.83 ± 1.92 | compared to N2 C, *p* = 0.0305 |
|  | *mir-249* | Exercise | 60/60 | 21.97 ± 1.58 | compared to *mir-249* C, *p* < 0.0001 |

Mean lifespan and statistical analysis were determined by OASIS2 platform, Kaplan-Meier curve was performed using GraphPad Prism 7.
Number of worms represents the number of dead worms scored relative to total number of worms initially started with. The difference in number is indicative of censored worms.

**Table S3.**
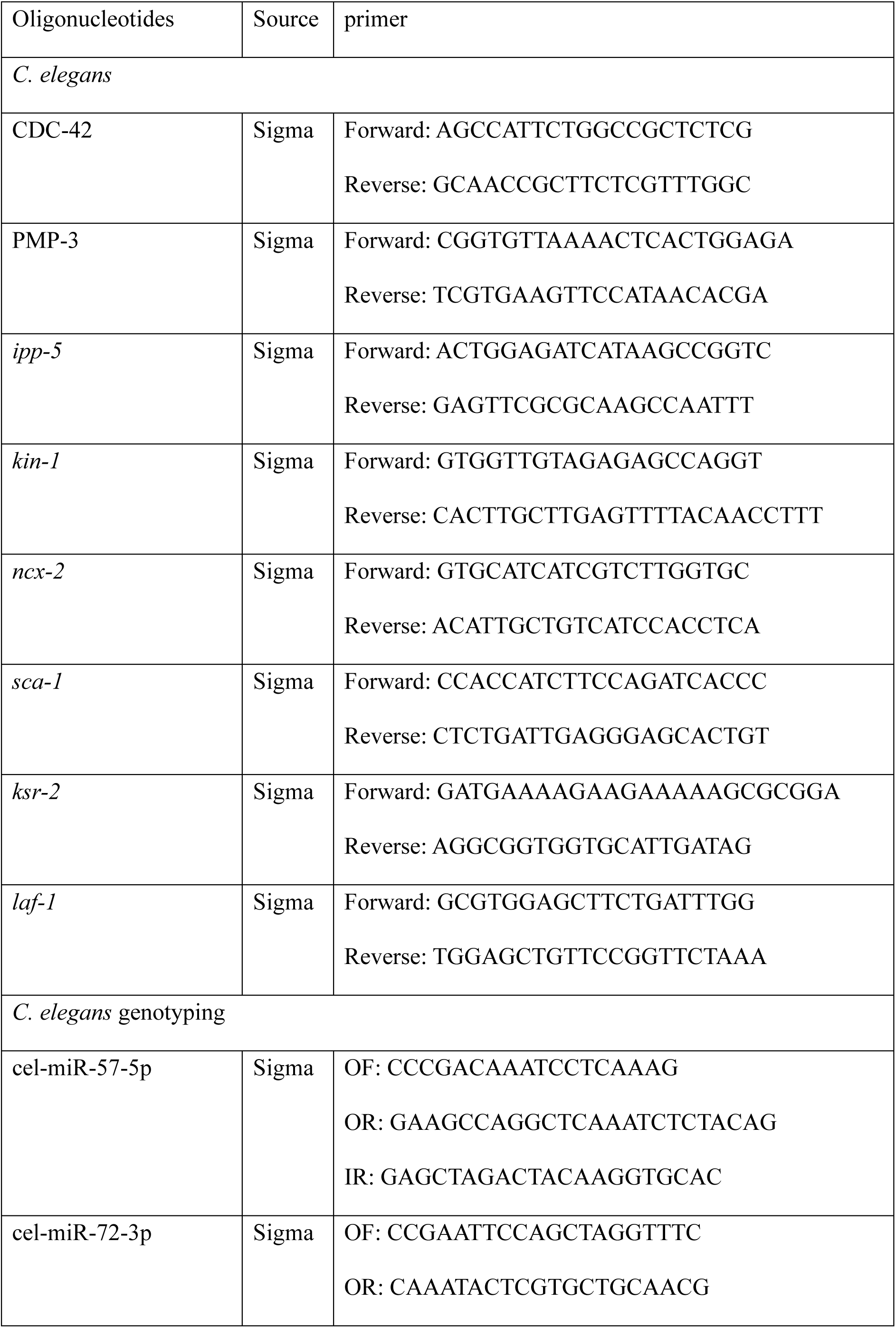
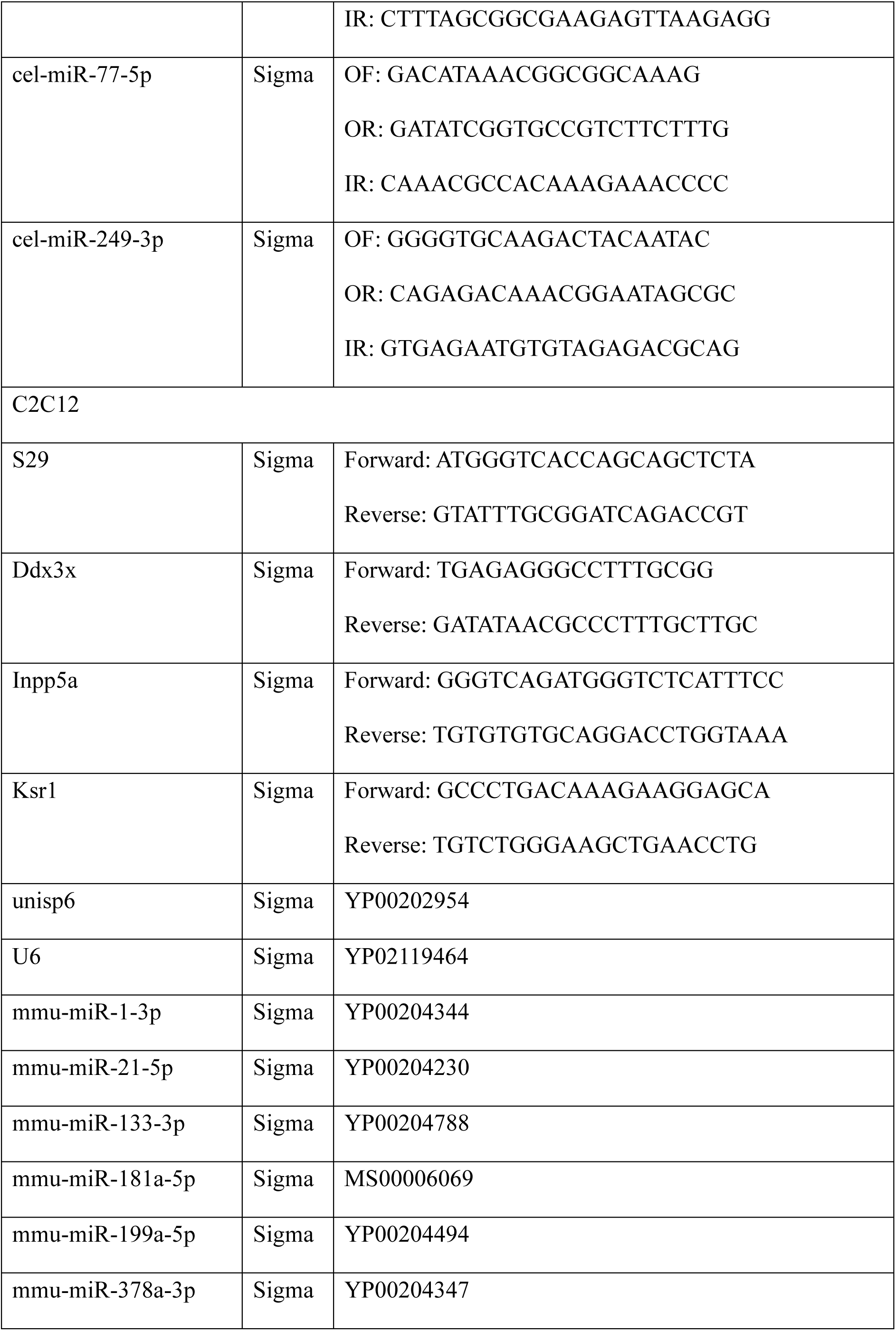
qPCR and genotyping primers.

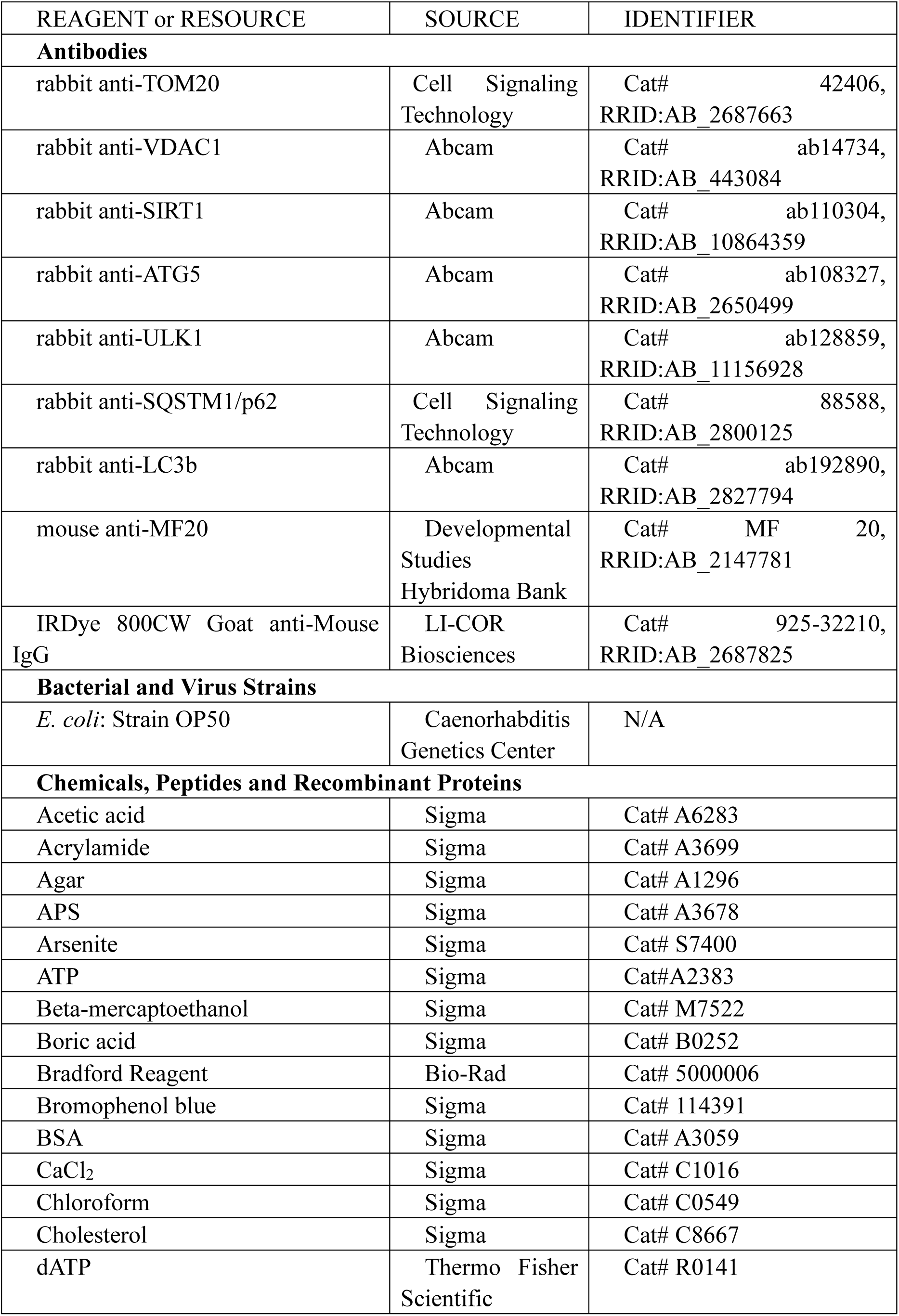
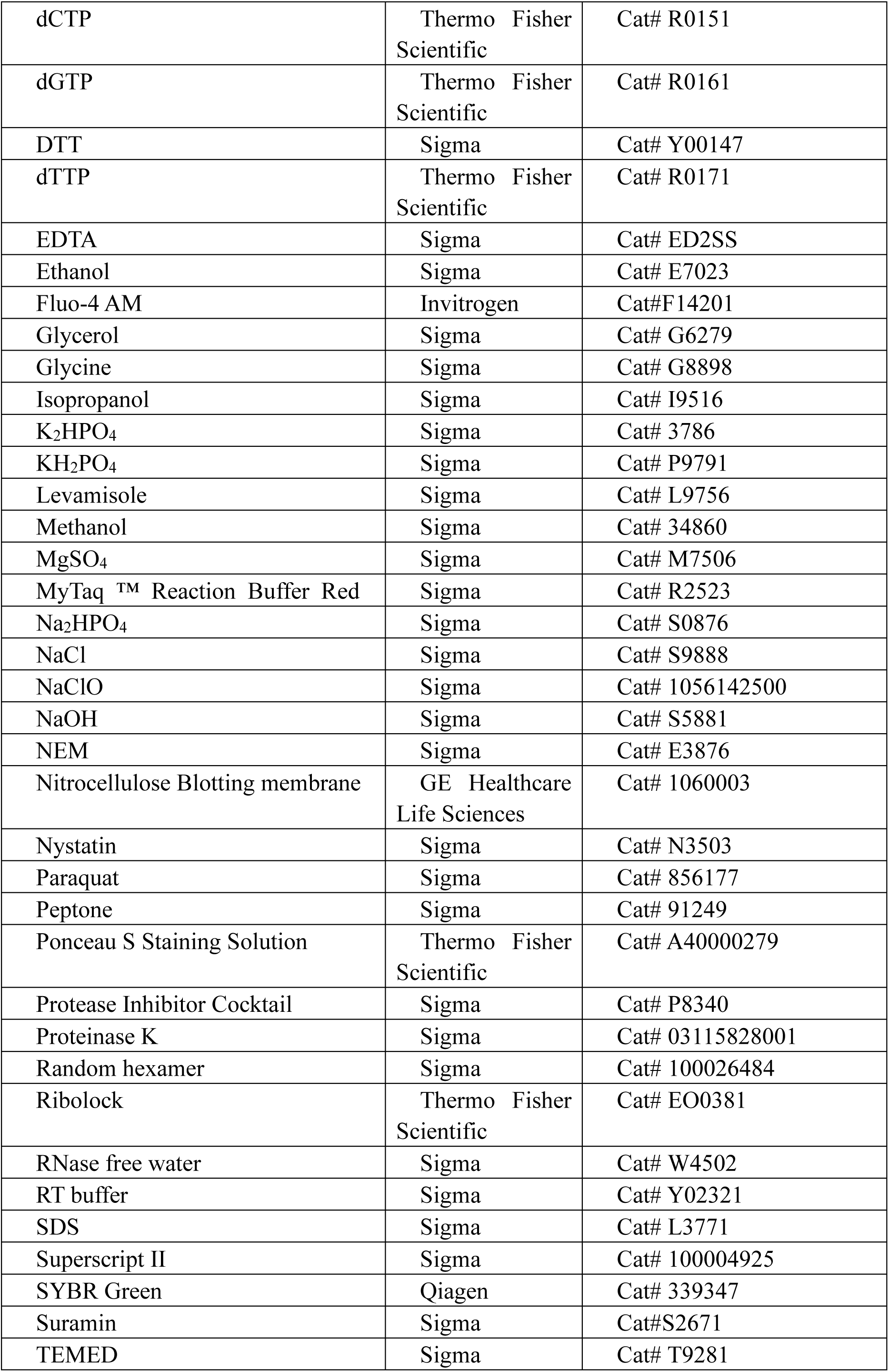
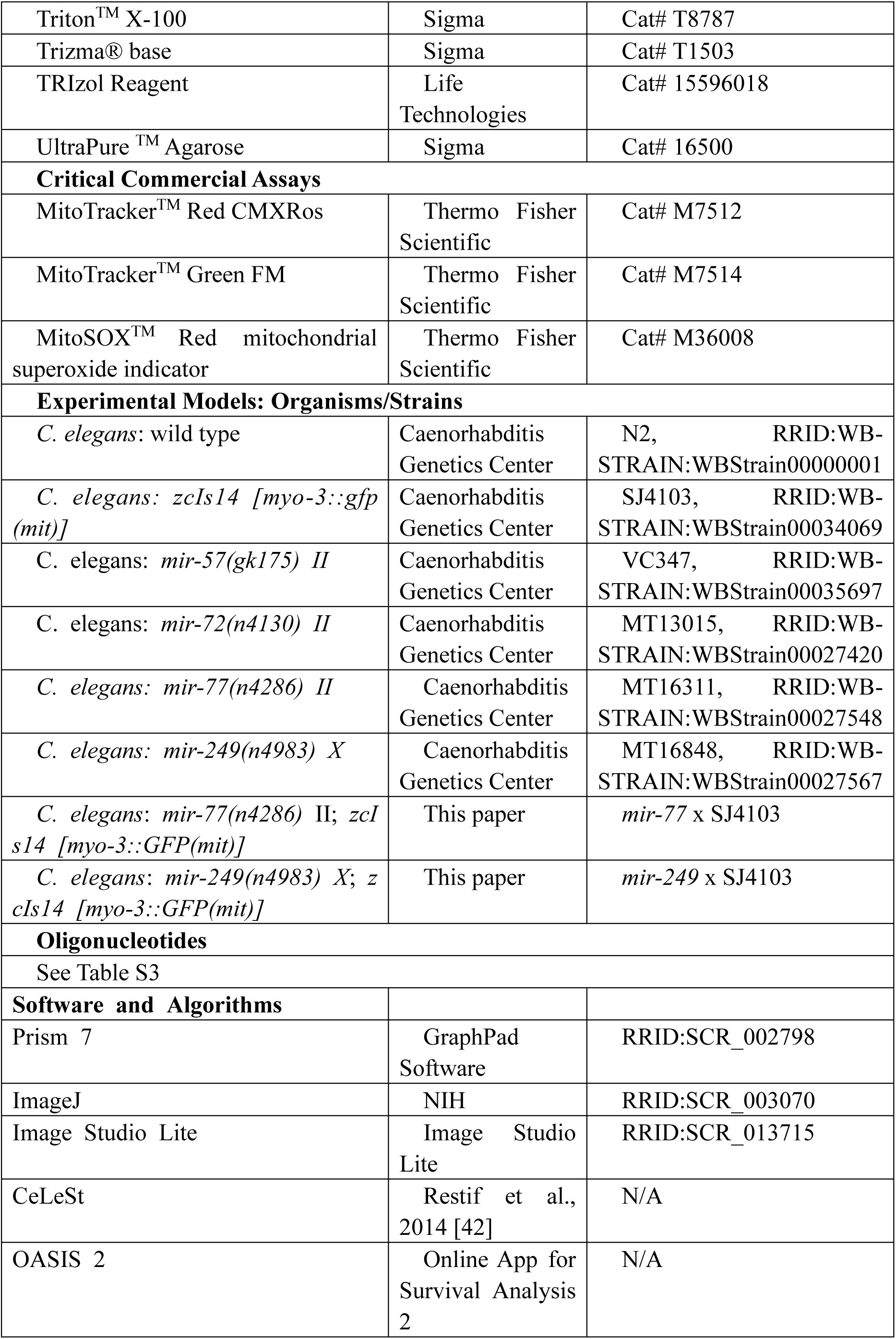
Reagents and Resources:

